# iPSC-derived Cancer Organoids Recapitulate Genomic and Phenotypic Alterations of c-*met*-mutated Hereditary Kidney Cancer

**DOI:** 10.1101/518456

**Authors:** Jin Wook Hwang, Christophe Desterke, Olivier Féraud, Stephane Richard, Sophie Ferlicot, Virginie Verkarre, Jean Jacques Patard, Julien Loisel-Duwattez, Adlen Foudi, Frank Griscelli, Annelise Bennaceur-Griscelli, Ali G Turhan

## Abstract

Hereditary cancers with cancer-predisposing mutations represent unique models of human oncogenesis as a driving oncogenic event is present in germline, exposing the healthy member of a family to the occurrence of cancer. The study of the secondary events in a tissue-specific manner is now possible by the induced pluripotent stem cell (iPSC) technology offering the possibility to generate an unlimited source of cells that can be induced to differentiate towards a tissue at risk of malignant transformation. We report here for the first time, the generation of a c-*met*-mutated iPSC lines from the somatic cells of a patient with type 1 papillary renal cell carcinoma (PRCC). We demonstrate the feasibility of kidney differentiation with iPSC-derived organoids expressing markers of kidney progenitors with presence of tight junctions and brush borders in tubular structures at transmission electron microscopy. Importantly, c-*met*-mutated kidney organoids expressed PRCC markers both *in vitro* and *in vivo* in NSG mice. Gene expression profiling of c-*met*-mutated iPSC-derived organoid structures showed striking molecular similarities with signatures found in a large cohort of PRCC patient samples and identified 11 common genes. Among these, *BHLHE40 and KDM4C*, well-known factors involved in PRCC pathogenesis, were expressed in c-*met*-mutated kidney organoids. This analysis applied to primary cancers with and without c*-met* mutation showed overexpression of the BHLHE40 and KDM4C only in the c-*met*-mutated PRCC tumors, as predicted by c-*met*-mutated organoid transcriptome. These data represent therefore the first proof of concept of the generation of “renal carcinoma in a dish” model using c-*met*-mutated iPSC-derived organoids, opening new perspectives for discovery of novel potentially predictive disease markers and novel drugs for future precision medicine strategies.

## INTRODUCTION

Hereditary cancers are due to oncogenic mutations or deletions and represent a major challenge in terms of diagnosis, prognosis and prevention^1^. Recently, the development of the iPSC technology allowed these cancers to be modeled using patient derived pluripotent stem cells^2–4^. It has been shown that this strategy can be used for modeling BRCA1-mutated breast cancer^5^ and Li-Fraumeni syndrome with TP53 deletions^3^. The major problem regarding the modeling of cancer using iPSC is that the effects of oncogenic mutations may appear only in the tissue in which the cancer develops^6^. Indeed, in a model of pancreatic cancer using iPSC, cancer phenotype appeared only when the specification of the endoderm had been achieved^6^. Similarly, the use of drug targeting strategies requires the generation of tissue-specific organoids, a technology which has been achieved in many tissues using either primary cells or pluripotent stem cells^7^. However, the feasibility and clinical use of iPSC-derived cancer organoids harboring a hereditary oncogenic mutation has not been shown so far. For this purpose, we have generated iPSC lines from a patient with hereditary c-*met*-mutated PRCC. Kidney organoids from the patient-specific iPSC were developed using a 3D *in vitro* culture system for immunochemistry, transmission electron microscopy and transcriptome studies. The results reported here show the possibility of generating kidney organoids with phenotypic PRCC markers using an in vitro, two-step 3D culture system and in vivo after injection into NSG mice. Strikingly, transcriptome analysis of c-*met*-mutated iPSC-derived renal progenitor cells revealed a gene profiling pattern similar to that reported in c-*met*-mutated primary human PRCC In addition, the markers that we have discovered in c-*met*-mutated kidney organoids are also overexpressed only in primary tumors of c-*met*-mutated PRCC, validating the feasibility of generating unlimited sources of cancer organoids for future drug screening strategies.

## RESULTS

### Patients

The propositus is a patient in whom the diagnosis of the c-*met* mutation was performed during a genetic study realized in his family 5 additional patients with (n=2) and without (n=3) c-*met* mutated PRCC have been analyzed in the study.

### Generation and characterization of c-*met*-mutated iPSC

Cellular programming was performed using the patient’s bone marrow mononuclear cells after her informed consent and the approval of Inserm ethical committee. c-*met*-mutated iPSC was generated using Sendaï virus mediated gene transfer of the four Yamanaka factors (**Supplementary Fig. 1a**). As a control we used iPSC generated from bone marrow mononuclear cells of a normal donor. Flow cytometric analysis demonstrated that the majority of c-*met*-mutated iPSC as well as control iPSC coexpressed SSEA4 and TRA-1-60 (91.3% and 89.8% respectively) markers (**Supplementary Fig. 1b and 1c**) as expected for human pluripotent stem cells. *In vivo* teratoma induction assay in NSG recipient mice revealed differentiation towards all three germ layers (**Supplementary Fig. 1d**) such as neural rosettes (ectoderm), immature cartilage (mesoderm), and glandular epithelia (endoderm). The same results were obtained using control iPSCs (**Supplementary Fig. 1e**). These data demonstrated the successful generation of fully reprogrammed c-*met*-mutated iPSC using this technology **Generation of c-*met*-mutated iPSC aggregates (iPSC-A)**.

To demonstrate that c-*met*-mutated iPSC can lead to formation of kidney organoids spontaneously, we cultured iPSC-A under ultra-low attachment conditions (**Supplementary Fig. 3**). As shown in **Supplementary Fig. 3c**, c-*met*-mutated iPSC-A cultured for up to 6 days were evaluated for the expression of kidney related markers SIX2 (SIX homeobox 2, a marker of kidney progenitors) and PODXL (podocalyxin like, a marker of glomerular podocytes) as described^9,10^. Immunochemistry analyses showed an the detection of both SIX2- and PODXL-positive cells at day 3 and day 6 (**Supplementary Fig. 3b,c**). c-*met*-mutated iPSC-A were then further characterized using immunocytochemistry for expression of phospho-Met and Brachyury. These experiments demonstrated that both control and c-*met*-mutated iPSC-A contained cells expressing phosphorylated c-Met protein (phospho-Met) and expressing Brachyury at 1 and 6 days (**Supplementary Fig. 2c,d**). Of note, the level of phospho-Met expression was higher in both control and c-*met*-mutated iPSC-A culture as compared to the monolayer counterparts (**Supplementary Fig. 2b**). These data demonstrate that aggregate formation strongly favors the emergence of double-positive SIX2 and phospho-Met expressing kidney progenitors from both control and c-*met*-mutated iPSC-A.

### Evaluation of a 3D culture system for induction of kidney differentiation from c-*met*-mutated iPSC

We performed a 3D culture system protocol to induce kidney differentiation for 12 days. (**Supplementary Fig. 3d,e**). 3D culture system was compared with the monolayer culture (day 6) of both control and c-*met*-mutated iPSCs to evaluate the efficiency of differentiation into kidney at day 12 (**Supplementary Fig. 4a**). The expression of kidney markers von Hippel-Lindau (VHL) and PODXL was determined by immunocytochemistry. As seen in **Supplementary Fig. 4b**, the 3D culture system outperformed the monolayer conditions in terms of level of VHL and PODXL expression. When only a weak expression of both proteins is observed in cells cultured in monolayers, a strong signal is noticed in cells grown in 3D cultures in both groups (**Supplementary Fig. 4c**). This result indicates that 3D cultures surpassed monolayers as they robustly enabled the generation of VHL- and PODXL-positive renal cells.

### Characterization of kidney organoids

To investigate whether we can generate kidney organoids from the iPSC-A using our 3D culture system, control iPSC-A as well as c-*met*-mutated iPSC-A were plated on 24- or 96-well Geltrex-coated dishes or 8-well culture chambers and cultured for another 6 days (**Fig. 1a and Supplementary Fig. 5a**). At 12 days immunocytochemistry was performed using antibodies against phalloidin (a cytoskeleton marker) and PODXL. After 12 days of differentiation, we obtained two types of kidney organoids based on their 3D shape, designed as “cup-like organoid” type (**Supplementary Fig. 5b,d,f**) and a “fusion organoid” type (**Supplementary Fig. 5c,e,g**). Immunocytochemistry revealed that in the “cup-like organoids” with cavitation, PODXL expression was markedly detected in the center and periphery of the organoids whereas phalloidin expression was restricted to the periphery (**Supplementary Fig. 5b,d**). In the “fusion organoid” type structures we detected strong expression of both PODXL and phalloidin throughout (**Supplementary Fig. 5c,e**). These results indicate that renal differentiation of c-*met*-mutated iPSC could be assessed within 2 weeks with the formation of specific kidney organoids expressing PODXL glomerular marker. We then performed kidney organoids from both normal and c-met-mutated iPSC to compare the expression of glomerular or tubular markers at different time points during this differentiation process. As seen in **Fig. 1**, our organoids markedly expressed the glomerulus-specific marker Nephrin, the endothelial marker CD31 and the lotus tetragonolobus lectin (LTL) a marker of proximal tubule epithelial cells (**Fig. 1b,e**), as expected phospho-Met was strongly expressed in kidney organoids with c-*met* mutation (**Fig. 3c and Fig. 5h**). Interestingly glomerulus and tubulin structure in normal kidney organoids seems to be spatially organized (**Fig. 1b,d**), as shown by staining distribution, and shape of the organoids than the c-*met* mutated kidney organoids (**Fig. 1e,g**).

**Fig. 1.**
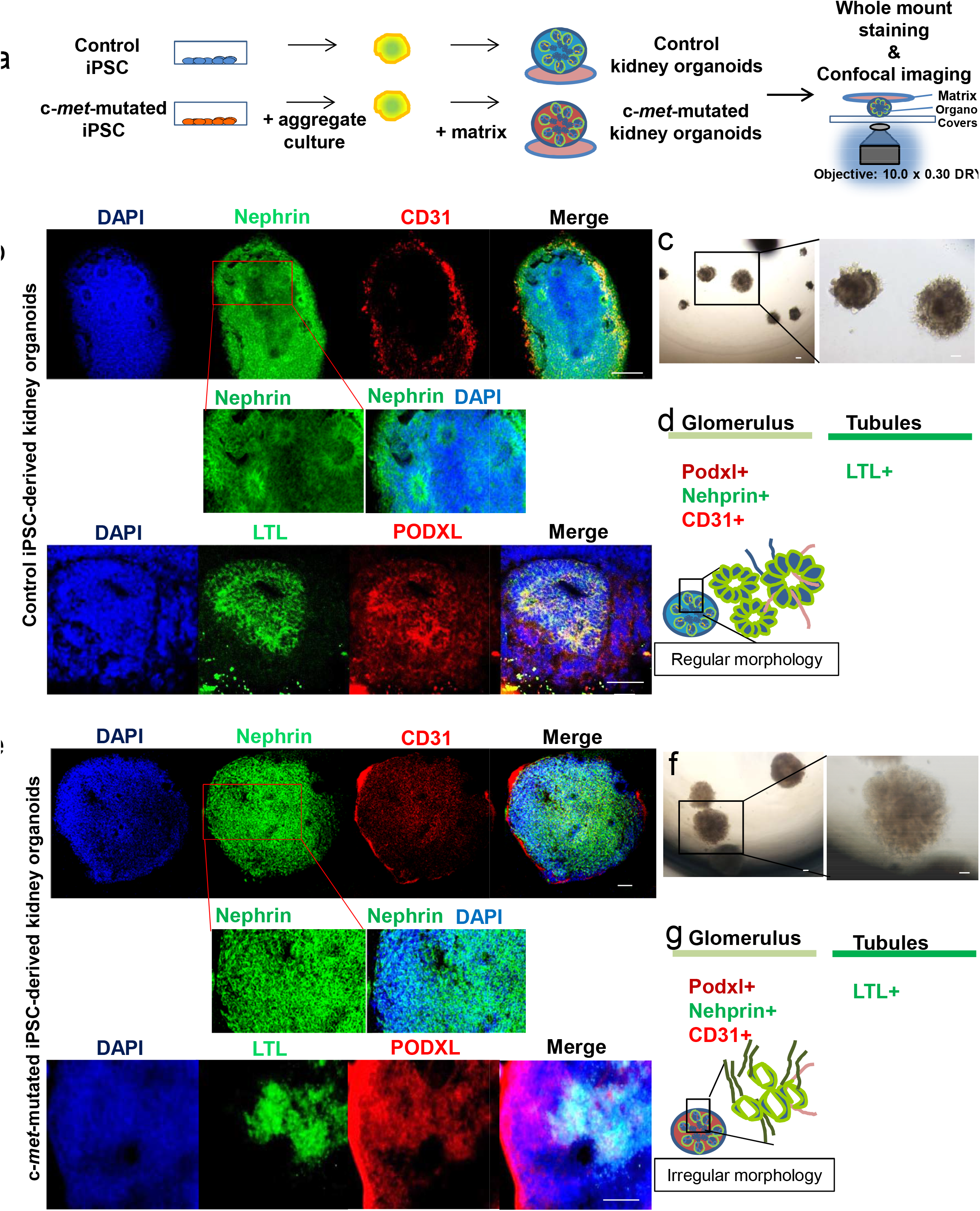
Kidney organoids from control or c-*met*-mutated iPSC in 3D culture. **a,** Schematic representation of the generation of iPSC-derived kidney organoids from control or c-*met*-mutated iPSC. **b,** Confocal analysis and whole-mount staining for Nephrin, CD31, LTL, PODXL, and DAPI in normal iPSC-derived kidney organoids showing nephron vesicles and absence of phospho-Met staining. Scale bar : 100 μm. **c,** Optical image of control iPSC-derived kidney organoids at day 12 showing the formation of tubule structures. Scale bar : 100 μm. **d,** Schematic representation of glomerulus and tubule markers with nephron formation vesicle, nephron structure and whole-mount staining for Nephrin. **e,** Confocal analysis and whole-mount staining for Nephrin, CD31, LTL, PODXL and DAPI in c-*met*-mutated iPSC-derived kidney organoids, Scale bar : 100 μm. **f,** Optical image of c-*met*-mutated iPSC-derived kidney organoids at day 12, Scale bar : 100 μm. **g,** Schematic representation of glomerules and tubules with appropriate markers showing nephron vesicles and nephron structures. The c-*met*-mutated organoid-derived vesicles **(e)** present an irregular morphology as compared to controls **(b)**.

### Structural and functional evaluation of kidney organoids

Spatiotemporal organization and ultrastructure of kidney organoids were analyzed by performing transmission electron microscopy (TEM). As can be seen in **Fig. 2a**, these experiments clearly showed ultrastructures corresponding to the generation of tight junctions (**Fig. 2a**) and brush border-like structures as can be expected to be seen in proximal tubules (**Fig. 2a**). Interestingly, the brush borders detected by TEM in c-*met*-mutated iPSC-derived kidney organoids were well developed as compared to brush borders from control iPSC-derived kidney organoids (**Fig. 2a,d**). In the same sections used for TEM, these structures have been found to express PODXL and LTL, confirming the presence of glomerular and tubular cells derived from c-*met*-mutated iPSC or control iPSC (**Fig. 2c,f**). TEM analyses performed at day +12 of the differentiation, showed a mature epithelial ultrastructure (**Fig. 4b,c**). Podocyte-like cells with foot processes were observed in glomeruli structures (**Fig. 2a,d**). Interestingly, large amounts of glycogen granules were observed in c-*met*-mutated iPSC-derived kidney organoids (**Fig. 4c,f,g**). In all iPSC-derived kidney organoids, tubules comprised mitochondria-rich cells with brush border-like structures (**Fig. 2a and Supplementary Fig. 6d**). We then analyzed the functionality of proximal tubule structures by using a dextran uptake assay in organoids at day 12. These assays demonstrated the selective uptake of dextran-Alexa488 from the media after 48 h of exposure in kidney organoids (**Fig. 2g,h**). These findings demonstrated that the organoid structures obtained in our experiments corresponded to kidney cells generated *in vitro* from c-*met*-mutated and control iPSC.

**Fig. 2.**
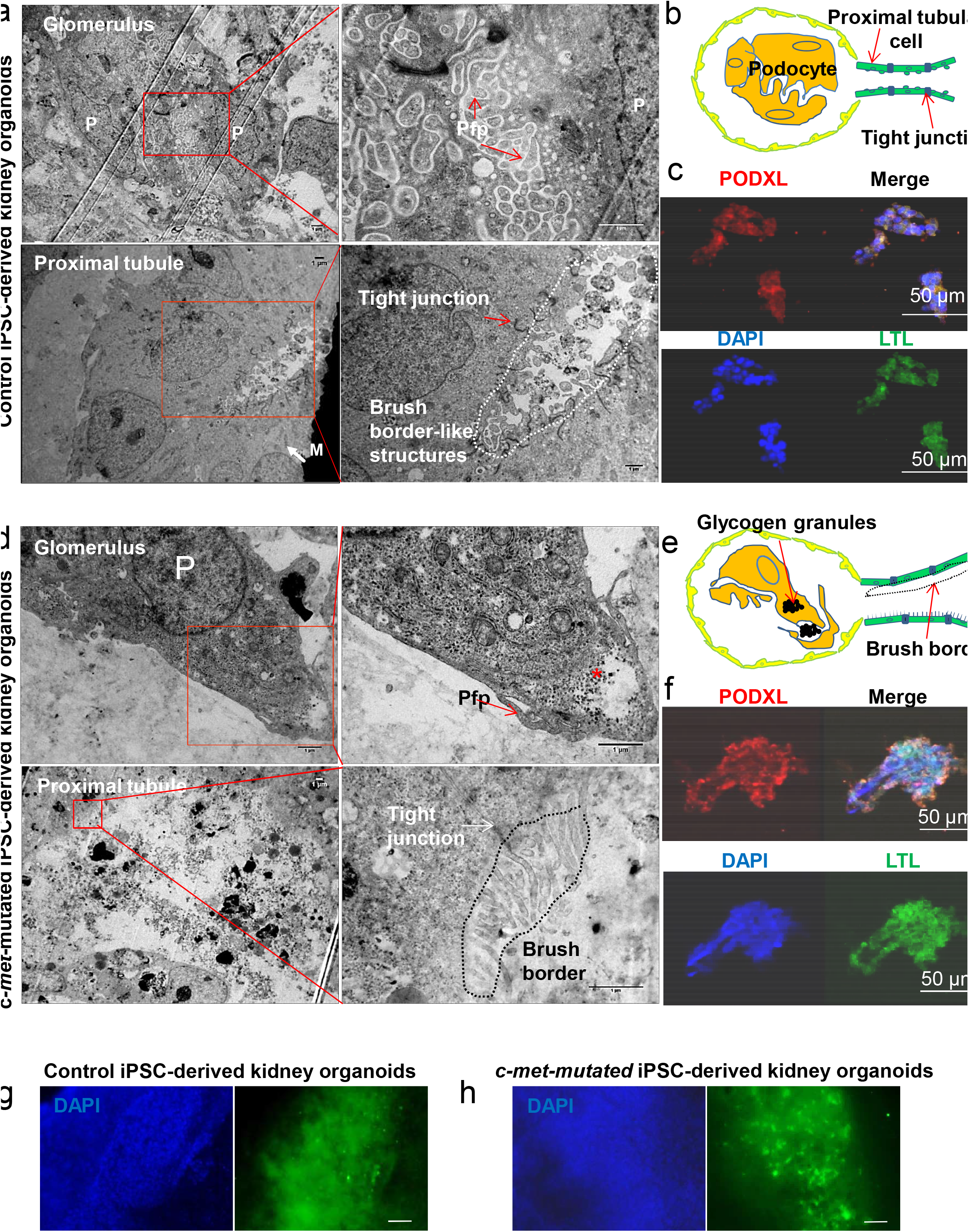
Ultrastructure and immunohistochemistry analyses by confocal microscopy. **a-c:** Representative electron microscopy images of glomeruli and tubules of control and c-*met*-mutated iPSC-derived kidney organoids showing podocyte-like cells (P), podocyte-like foot process (Pfp), mitochondria (M) and brush borders. The immunohistochemistry analysis in paraffin cuts reveals both PODXL and LTL positivity in organoids. Scale bar : 50 μm. **d-f:** Representative electron microscopy images glomeruli and tubules of c-*met*-mutated iPSC-derived kidney organoids revealing podocyte-like cells (P), podocyte-like foot process (Pfp), tight junctions and typical brush border structures. Immunohistochemistry analysis in paraffin cuts by confocal imaging showed organoid structures co-expressing PODXL-and LTL. Scale bar : 50 μm. As compared to control iPSC-derived organoids, the presence of glycogen granules(*) were noted **(d) g, h:** Evaluation of functional analysis of proximal tubules structures in kidney organoids by using Dextran uptake assays. Organoids were incubated for 48 hours in the presence of Dextran-Alexa 488 followed by analysis using wide-field microscopy. Dextran uptake was seen in normal **(g)** as well as in c-*met*-mutated **(h)** organoids with a more intense uptake in the latter structures **(see text)**. Scale bar : 100 μm.

By using kidney organoid model, we asked whether biomarkers expressed in kidney cancer, could be expressed in c-*met*-mutated iPSC-derived kidney organoids. For this purpose, we performed immunostaining using cytokeratin 7 (Cy7)^11^, cubilin^12^ well-established markers of type 1 PRCC (**Fig. 3b,c**). As can be seen in **Fig. 3b**, cytokeratin 7 and cubilin expression were detected in c-*met*-mutated iPSC-derived kidney organoids (**Fig. 3b**) whereas transcription factor for immunoglobulin heavy chain enhancer 3 (TFE3)^13^ expression, a known kidney cancer marker, was expressed at similar level to that observed in control iPSC-derived kidney organoids (**Fig. 3b,c**). As compared with control iPSC-derived kidney organoids, phospho-Met signal was strongly detected in all c-*met*-mutated iPSC-derived kidney organoids (**Fig. 3b,c**).

**Fig. 3.**
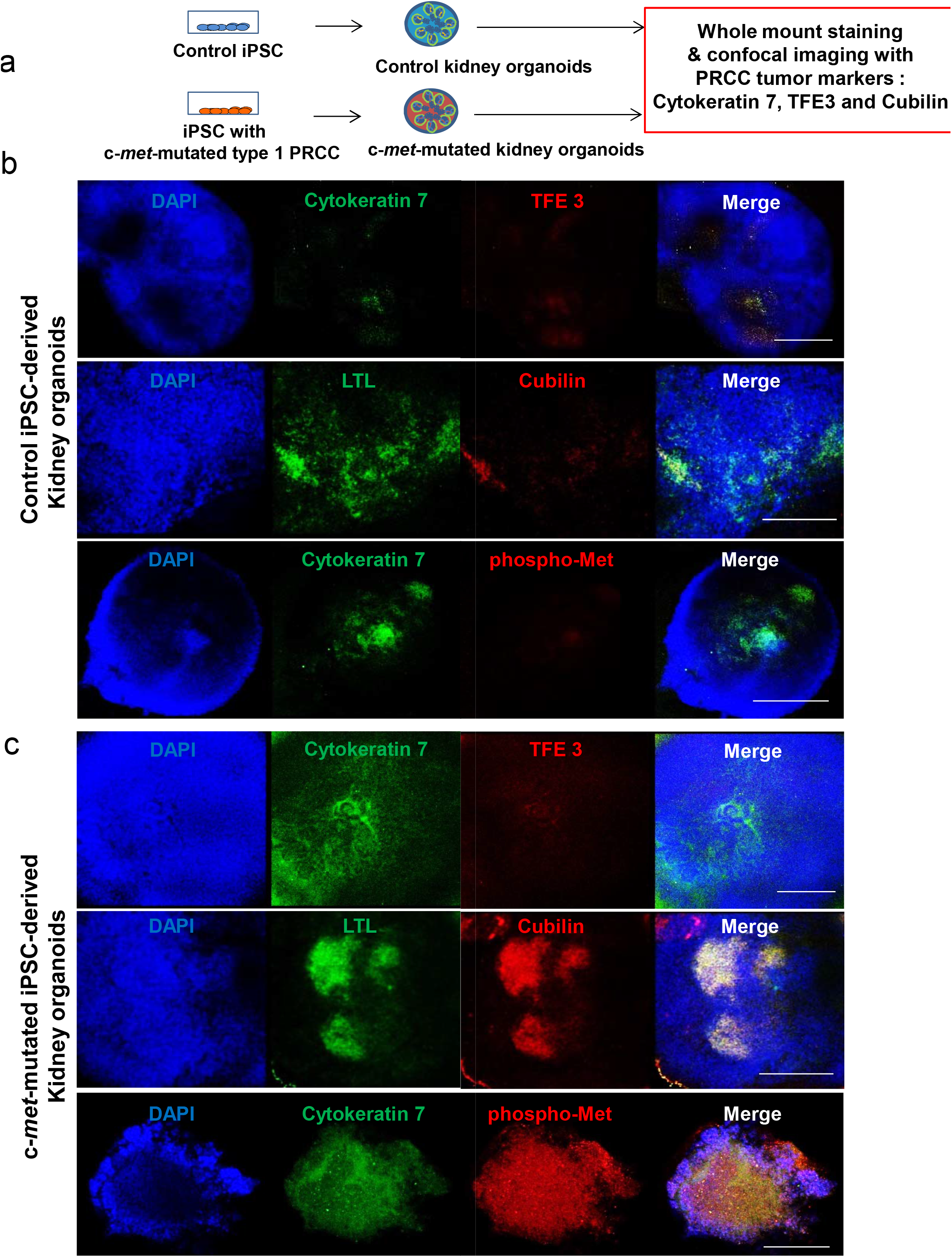
Characterization of c-*met*-mutated iPSC-derived kidney organoids. **a,** Schematic representation of the generation of control or c-*met*-mutated iPSC-derived kidney organoids and characterization of clinicopathological cancer markers cytokeratin 7, TFE3 and Cubilin. **b,** Whole-mount staining for Cytokeratin 7, TFE3, LTL, Cubilin and DAPI in control iPSC-derived kidney organoids, Scale bar : 500 μm. **c,** Whole-mount staining for Cyto-keratin 7, TFE3, LTL, Cubilin and DAPI in c-*met*-mutated iPSC-derived kidney organoids. Scale bar : 500 μm.

### Kidney capsule transplantation experiments of c-*met*-mutated kidney organoids

To determine if c-*met*-mutated iPSC derived organoids can recapitulate the features observed *in vivo*, we transplanted them under the kidney capsule of NSG mice. To this end, we transplanted kidney organoids derived from control and c-*met*-mutated iPSC into different groups of mice. At 4 weeks post-transplant, mice were sacrificed, and harvested structures were subjected to histopathological and immunocytochemistry analyses. We observed that c-*met*-mutated iPSC generated larger teratoma-like tumors compared to controls (**Supplementary Fig. 9b,c**). Histopathological analyses using H&E staining revealed the presence of immature cartilage in tumors of c-*met*-mutated iPSC group (**Supplementary Fig. 9f**). Moreover, kidney organoids with aggressive form were observed in the cross-sections of c-*met*-mutated kidney organoids (**Supplementary Fig. 9h,g**). In contrast, control showed a well-preserved form of kidney organoids (**Supplementary Fig. 9d**). Immunostaining was performed to determine glomerular or tubular origin of such structures. Indeed, the expression of Nephrin and CD31 (**Supplementary Fig. 9e,i**) in both groups demonstrated the presence of podocytes and endothelial cells in kidney-like structures. Notably, expression of PRCC markers cytokeratin 7 and TFE3 (**Supplementary Fig. 9e,i**) were found markedly increased in c-*met*-mutated kidney organoids-derived tissue as compared to controls. Altogether, these data suggest that a PRCC-like phenotype can be recapitulated *in vivo* using c-*met*-mutated iPSC-derived organoids.

### Drug toxicity assays

To test whether our organoids could be used to study kidney toxicity *in vitro*, we tested two different drugs on our c-*met*-mutated iPSCs-derived kidney organoids. After 12 days of differentiation, organoids were treated for 3 days with two doses of Sunitinib (either 200 μg/well or 5000 μg/well), a commonly used tyrosine kinase inhibitor in PRCC therapy. Upon treatment, we analyzed the presence of the Kidney Injury Molecule-1 (KIM-1), a biomarker upregulated in the proximal tubules following acute kidney injury. As seen in **Supplementary Fig. 8h,i**, KIM-1 was detected at low level in organoids treated with low dose of sunitinib whereas a higher dose of Sunitinib resulted in higher levels of KIM-1 (**Supplementary Fig. 8h,i**). Similarly, c-*met*-mutated iPSC-derived kidney organoids treated with cisplatin harbored KIM-1 upregulation albeit to lesser degree than that observed in control iPSC-derived kidney organoids. These data showed that c-*met*-mutated iPSC-derived kidney organoids could be used to test the nephrotoxicity and perhaps evaluate the therapeutic efficacy of novel candidate drugs *in vitro*.

### c-*met*-mutated iPSC aggregates reproduce molecular features of human PRCC

In order to determine the gene expression pattern of c-*met*-mutated iPSC, we performed transcriptome analysis on both c-*met*-mutated iPSC and c-*met*-mutated iPSC-A. Supervised analysis between the 2 classes of culture c-*met*-mutated iPSC versus c-*met*-mutated iPSC-A by ranking products algorithm enabled us to identify 196 differentially expressed gene probes (**Supplementary Table 2**). Of which, 148 were found to be down-regulated in c-*met*-mutated iPSC-A as compared to c-*met*-mutated iPSC. A small fraction of them (n=48) were found to be up-regulated in c-*met*-mutated iPSC-A as compared to c-*met*-mutated iPSC (**Fig. 5a and Supplementary Table 2**). In parallel, we investigated analysis of TCGA consortium Next Generation Sequencing dataset of PRCC tumor samples from the most recent cohort study of transcriptome with mutation sequence status that was available^14^. This allowed us to stratify the transcriptome of PRCC by their c-*met* mutation status: 23 mutations were found carried by 21 unique patients (**Supplementary Table 4**). Machine learning supervised by c-*met* status performed on PRCC RNA-seq samples allowed to characterize 1333 predictive genes with a minimum error of misclassification (data not shown). Meta-analysis between c-*met*-mutated iPSC signature and PRCC signature revealed a significant enrichment of c-*met*-mutated iPSC profile as predictive of c-*met*-mutated PRCC tumor status (Fold of enrichment: 5.68; p-value<2.2E-16, **Fig. 5b**). Unsupervised principal component analysis performed with *met*a-analysis intersection genes (77 genes, **Supplementary Table 3**) on the transcriptome of PRCC tumor patients confirmed a significant stratification of these tumor samples taking into account their c-*met* mutation status (p-value=2.25E-10, **Fig. 5c**). Expression heat map performed on PRCC tumors samples with the 77 *met*a-analysis intersection genes (**Fig. 5d**) revealed a distinct expression pattern of these genes in PRCC patients that carried c-*met* mutations as compared to others; also the proportion of up regulated and down regulated genes is even in this subgroup of patients. These results suggest that the gene expression profile of c-*met*-mutated iPSC aggregates is an efficient model predictor of PRCC tumor stratification taking into account their c-*met* mutation status. In order to understand the influence of c-*met*-mutated iPSC-A signature study in the PRCC, a protein-protein interaction (PPI) network was built with extraction of the protein interactions from innateDB database concerning the 77 genes found at the intersection of the meta-analysis (**Supplementary Table 3**). This interaction network analysis allowed to build a principal network comprising 65 seeds, 1713 nodes and 2204 edges (**Fig. 5e**). Analysis of topology network importance revealed central connectivity of key molecules such as EGR1 and other stem cell related molecules (EZH2, NOTCH2, GLI3) (red nodes on **Fig. 5e**). Functional inference performed on this interaction network performed with projection of KEGG pathway database revealed significant implication of genes already involved in renal carcinoma (green nodes on **Fig. 5e**, and green bar on **Fig. 5f**). Among these 26 kidney cancer genes, HGF which is the ligand of c-*met* was predicted to be connected in this network (**Fig. 5e**). Fold-change concordance analysis between c-*met*-mutated iPSC-A model and PRCC tumors revealed 11 upregulated markers with c-*met* mutation status (**Fig. 5g**). Among these eleven genes some were found with high connectivity on the previous interaction network such as KDM4C, which is implicated in chromosomal aberrations found in tumors and BHLHE40 an important regulator of circadian rhythms (ARNTL1 partner) and cell differentiation (**Fig. 5g**). This network analysis confirmed by inference that the use of iPSC-aggregates gene signature during *met*a-analysis can efficiently predict c*-met* status stratification of PRCC tumors. Furthermore, KDM4C expression was validated in our c-*met*-mutated iPSC-derived organoids as we observed a co-expression of KDM4C and phospho-Met in c-*met*-mutated iPSC-derived kidney organoids (**Fig. 5h**, lower panels). Control iPSC-derived kidney organoids exhibited low levels of KDM4C expression which did not overlap with that of phospho-Met (**Fig. 5h**, upper panels). These data demonstrate that c-*met*-mutated iPSC-derived kidney organoids recapitulate some molecular features of human PRCC and could be of major interest not only to model this hereditary cancer but also to further understand the molecular circuitry downstream of c-*met* during the progression of this hereditary disease.

### Transcriptome of c-*met*-mutated kidney organoids

To explore the genomic consequences of constitutionally active c*-met* signaling pathway in kidney cells and to explore the genomic circuitry activated in kidney organoids, we performed a global transcriptome analysis in the c-*met*-mutated kidney organoids compared to gene profile of controls. As seen in **Supplementary Table 1** and **Supplementary Figure 7**, 244 genes were found up-regulated and 71 down-regulated. Gene set enrichment analysis (GSEA) revealed an enrichment of targets involved in kidney development specifically in c-*met*-mutated kidney sample such as regulation of epithelial cell differentiation involved in kidney development (Normalized Enrichment Score (NES=+1.55, p<0.001, **Supplementary Fig. 7b**), renal tubule development (NES=+1.51, p<0.001, **Supplementary Fig. 7c**), and positive regulation of kidney development gene set (NES=+1.48, p<0.001, **Supplementary Fig. 7d**). Also, GSEA analysis revealed implication of pathophysiological process between these two kidney sample conditions with an enrichment of renal cell carcinoma (NES=+1.40, p<0.001, **Supplementary Fig. 7e**) and aging kidney gene sets (NES=+1.67, p<0.001, **Supplementary Fig. 7f**) specifically in c-*met*-mutated iPSC-derived kidney organoids compared to normal counterparts.

### Validation of the expression of markers detected in of c-*met*-mutated organoids in primary papillary renal carcinoma tumors

We next asked whether the candidate genes that we have found overexpressed in iPSC-derived c-*met*-mutated organoids were also expressed in primary papillary renal carcinoma. For this purpose, we analysed 5 primary PRCC, 2 from patients with c-*met* mutation (UPN4 and UPN5) and 3 from patients without c-*met*-mutation (UPN1, UPN2 and UPN3). One of the c-*met*-mutated PRCC sample (UPN5) is from the mother of our patient (**Supplementary Table 4**). Amongst the 11 candidate genes that we have previously identified, we focused on KDM4C and BHLHE40 genes which are known to be expressed in PRCC. To evaluate their level of expression, we performed immunohistochemistry in primary tumors derived from type 1 PRCC with and without and c*-met* mutation using specific antibodies. As seen in **Fig. 6**, KDM4C and BHLHE40 were markedly overexpressed in the c-*met*-mutated type 1 PRCC tumors (UPN4, UPN5, **Fig. 6d,e**) as compared to type 1 PRCC tumors without c-*met* mutation (UPN1, UPN2, UPN3, **Fig. 6a,b,c**). These results strongly suggested that the findings observed in c-*met*-mutated iPSC-derived kidney organoids are truly representative of tumor marker expression in the primary tumors from patients with PRCC.

## DISCUSSION

Germline mutations at the origin of family cancers represent a major challenge in oncology as there is no experimental models to study the future cancer development. c-*met*-mutated PRCC represents a major form of hereditary kidney cancer^15^ amongst kidney cancers which are the seventh most common malignancies in the United States^16^. PRCC includes tumors with indolent, multifocal presentation and solitary tumors with an aggressive, highly lethal phenotype. PRCC can be classified by morphology into types 1 and 2; the former being characterized by the presence of papillae and tubular structures covered with small cells containing basophilic cytoplasm and small, uniform oval nuclei. Type 1 disease is characterized by the activation of germline mutations of the c-*met* oncogene. We describe here the first model to our knowledge, of a c-*met*-mutated PRCC using iPSC-derived patient-specific kidney organoids. The first step of our experiments consisted on the design of experimental conditions allowing efficient and reproducible generation of kidney organoid differentiation. To this end, we have used 3D-culture of iPSC in the presence of E8 medium. As can be seen in **Fig. 1f and Supplementary Fig. 3e**, c-*met*-mutated iPSC can differentiate into kidney organoids spontaneously under 3D culture conditions. Indeed, the c-*met*-mutated iPSC-derived organoids were found to express kidney cell markers such as PODXL+, Nephrin+ and LTL+^10,17^.

Previous kidney organoid studies involved chemically defined protocols under monolayer culture conditions at the initial stage for human embryonic stem cells^18^ or human fibroblast-derived-iPSC ^18,19^because these conditions allow the control of long-term clonal growth and multilineage differentiation of the pluripotent cells. The large-scale expansion of these cells is relatively easy under monolayer culture conditions. Despite these advantages, such cultures lack cell–cell and cell–matrix interactions and fail to mimic cellular functions and signaling pathways occurring naturally in 3D culture conditions. One of the most efficient kidney organoid differentiation includes indeed a combination of monolayer and three-dimensional (3D) cultures^19^. The monolayer culture techniques require however, the use of various chemically defined factors to induce kidney differentiation. The method used here addressed these limitations by the occurrence of a self-organization during spontaneous development *in vitro* and circumvented the disadvantages of monolayer cultures requiring several stages of differentiation without induction of sufficient cell-cell and cell-matrix interactions. 3D culture conditions favoring these interactions generate the potential of differentiation into three germ layers via cellular microenvironment like embryoid bodies (EBs). Such EB culture conditions have been used to generate heart, kidney, and liver organoids^20^. Finally, we have found that 3D-culture conditions induced higher expression of VHL protein as compared to monolayer cultures creating a favorable condition for kidney organoid generation (**Supplementary Fig. 4b,c**). It is known that overexpression of VHL induces kidney cell differentiation through the integration of cell–cell and cell–matrix signaling in 786-O cells^21^. We have then used this 3D culture conditions to determine the possibility of generation of kidney organoids using c-*met*-mutated iPSC and control iPSC. As can be shown in **Supplementary Fig. 2b-d**, the comparison of monolayer cultures and 3D-cultures revealed the overexpression of phospho-Met in both c-*met*-mutated iPSC and control iPSC (**Supplementary Fig. 2d**). These phospho-Met overexpressing aggregates generated kidney organoids as demonstrated by the expression of PODXL in confocal microscopy experiments (**Supplementary Fig. 5b-g**) and that of nephrin and LTL (**Fig. 1**). As shown in **Supplementary Fig. 5b-g**, we have found that c-*met*-mutated iPSC aggregated spontaneously generating fusion structures undergoing cavitation. It is known that the cavitation process is essential for providing a free epithelial surface for the morphogenetic movement of epiblastic cells during the subsequent formation of a primitive streak^22^. This fusion could occur through the use CXCR4/CXCL12 axis, which is known to be essential for kidney vasculature development^23^. Indeed, immunostaining experiments showed the expression of CD31 in kidney organoids (**Fig. 1 b,e**). In order to demonstrate the efficient kidney organoid generation, we have used transmission electron microscopy experiments. These experiments confirmed, the appearance of kidney structures including glomeruli, podocytes, glomeruli-associated basement membranes and proximal tubule with typical brush border-like structures. Interestingly, in c-*met*-mutated iPSC-derived kidney organoids, high numbers of glycogen granules were observed as previously reported in primary tumors^24^ (**Fig. 4c,f,g**).

**Fig. 4.**
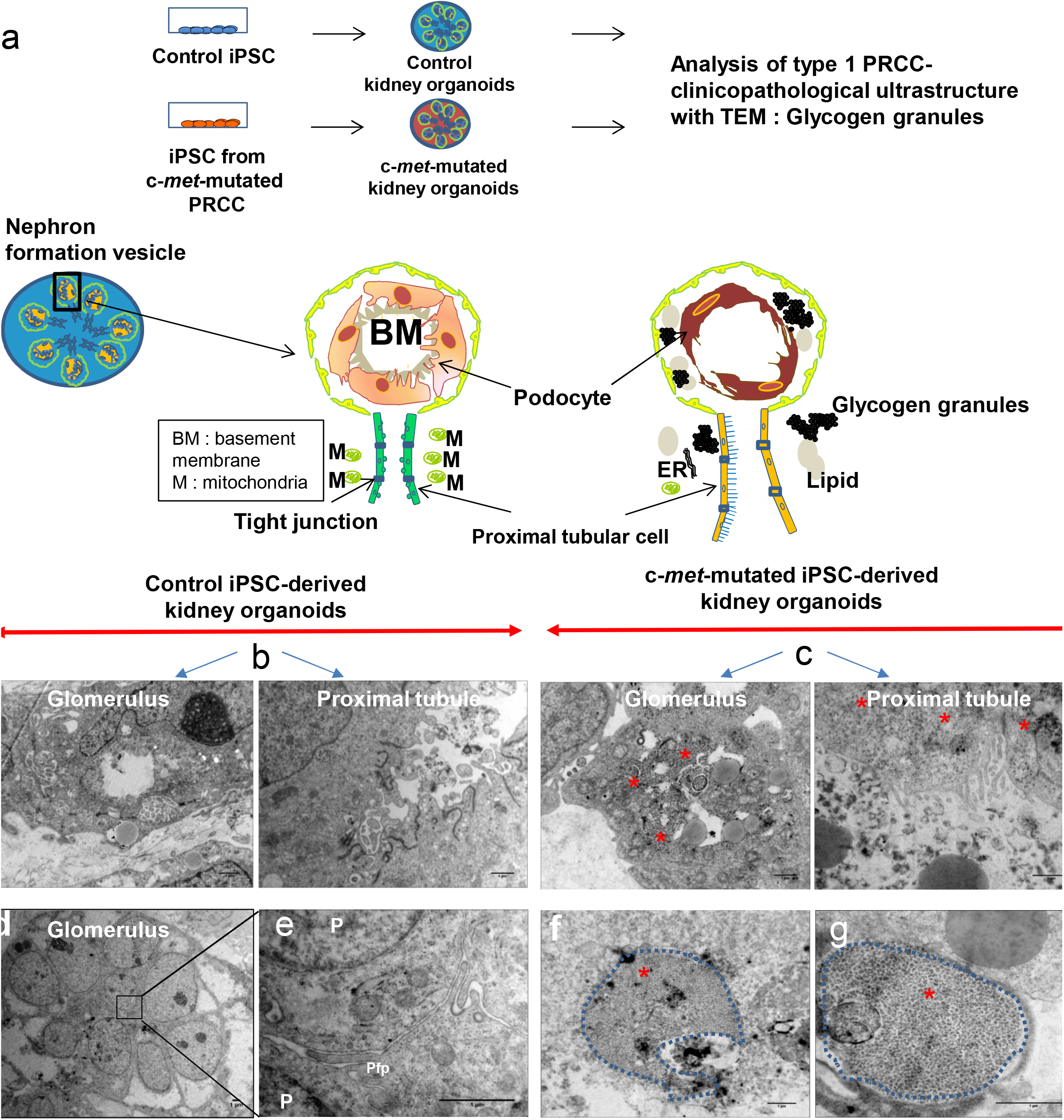
Ultrastructural analysis of kidney organoids derived from control or c-*met*-mutated iPSC. **a,** Schematic representation of the experimental protocol used. **b,c** Representative electron microscopy images of glomerulus and tubules structures of control or c-*met*-mutated iPSC-derived kidney organoids. **d,e** Representative electron microscopy images cytoplasm region of control iPSC-derived kidney organoids. **f,g,** Representative electron microscopy images cytoplasm region of c-*met*-mutated iPSC-derived kidney organoids, podocyte-like cells. Glycogen granules (*).

The next important question was to determine if some phenotypic cancer markers known to be expressed in human PRCC could be found in c-*met*-mutated iPSC-derived organoids. Cy7 overexpression was found in c-*met*-mutated iPSC-derived organoids as compared to control organoids, demonstrating the reproduction of a cancer phenotype in these structures (**Fig. 3c**). Moreover, in order to demonstrate the possible differential behavior *in vivo* of the c-*met*-mutated iPSC derived organoids, we have transplanted these cells as well as control organoids, under the kidney capsule of NSG mice using classical transplantation technology. 4 weeks after transplantation, mice were sacrificed and a necropsy was performed followed by pathological analysis. As can be seen in **Supplementary Fig. 9**, c-*met*-iPSC-derived organoids induced larger tumors as compared to controls and expressed kidney cancer markers TFE3 (transcription factor for immunoglobulin heavy chain enhancer 3)^13^ and Cy7^25^ demonstrating the generation in vivo cancer organoids from the c-*met* mutated iPSC.

We next asked whether a differential gene expression profiling can be obtained in c*-met* mutated iPSC and aggregates during the induction of aggregates as compared to control-iPSC-derived cells. This analysis allowed us to identify 196 differentially expressed genes, generating a clear transcriptome signature during the first stages of kidney organoid differentiation (**Fig. 5a**). Interestingly, 77 of these genes have also been found to be expressed specifically in c-*met*-mutated human kidney carcinoma (**Fig. 5b**) which appeared to be different from the signature observed in PRCC without c-*met* mutation. Most importantly, several of the 11 genes which have been identified as being overexpressed in our c-*met*-mutated iPSC, have also been overexpressed in primary human c-*met*-mutated PRCC such as KDM4C and BHLHE40 (**Fig. 5g**). KDM4C is a member of the Jumonji-domain 2 family encoding a de*met*hylase involved in chromosome segregation^26^. Alteration of KDM4C gene has been shown to occur in renal cell carcinoma ^27^. We next asked whether the overexpression of this gene, which has been reported in both our iPSC and primary human PRCC, could be reproduced in our c-*met*-mutated iPSC-derived kidney organoids. As can be seen in **Fig. 5h**, KDMC4 expression was clearly seen in phospho-Met-expressing c-*met*-mutated iPSC as compared to controls (**Fig. 5h**). Similarly, the basic helix-loop helix protein BHLHE40 known to be implicated in c-*met* mutated PRCC^14^ was found to be overexpressed in c-*met*-mutated iPSC derived organoids (**Fig. 5h**).

**Fig. 5.**
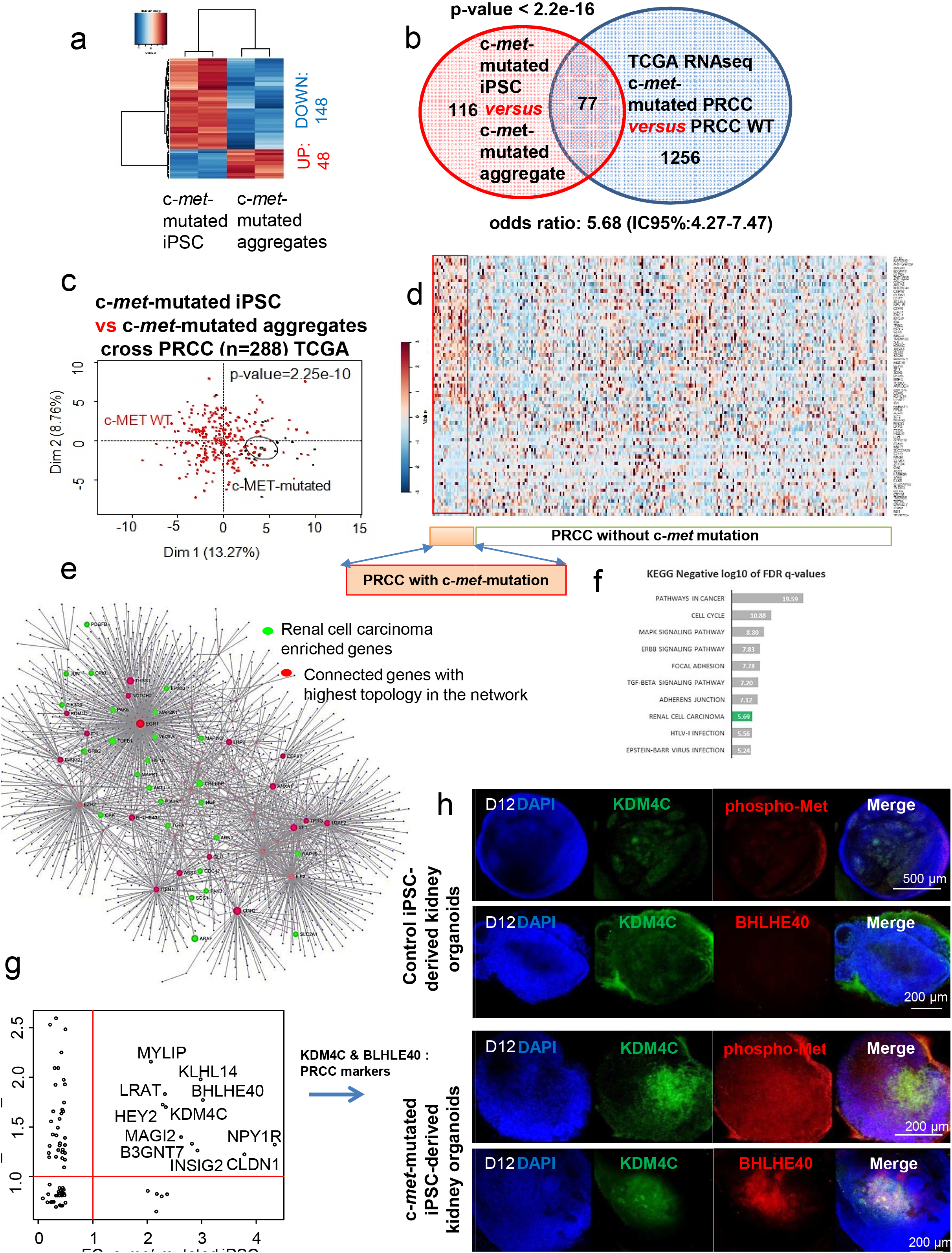
Transcriptome analyses of c-*met*-mutated iPSC aggregates. **a,** Expression heatmap (Euclidean distances) of differential expressed genes found between c-*met*-mutated iPSC (monolayer culture) and c-*met*-mutated iPSC aggregates. **b,** Venn diagram of the metaanalysis between c-*met*-mutated iPSC and PRCC patient analysis, p-value of the c-*met*-mutated iPSC signature in PRCC expression profile was calculated by hypergeometric test of Fisher. **c,** Principal component analysis performed with meta-analysis gene intersection (77 genes) on PRCC tumor samples (Z-scores RNAseq V2), p-value was calculated by correlation of the group discrimination of the first principal component. **d,** Expression heatmap performed with meta-analysis gene intersection (77 genes) on PRCC tumor samples. e, protein-protein interaction network built with the projection of 77 meta-analysis genes on innateDB interaction database: red genes represent connected genes with the highest topology in the network, green genes represent enriched gene during KEGG inference and related to the Renal cell carcinoma. **f,** Bar plot representing negative logarithm base 10 of False Discovery Rate (FDR) q-values found during functional inference of KEGG database on metaanalysis protein-protein interaction (PPI) network. **g,** Concordance scatterplot of fold-change (FC) found during meta-analysis of transcriptome. The x-axis shows FC of the genes found in c-*met*-mutated iPSC transcriptome study and the y-axis shows the FC of genes found in analysis of PRCC RNAseq study. The genes found to be expressed > 1 FC are shown by their gene symbols. **h,** Whole-mount staining for KDM4C (Lysine demethylase 4C), phospho-Met and DAPI in kidney organoids.

Finally, it was of importance to determine if the two markers that we have discovered through the analysis of c-Met-mutated iPSC could be truly representative of the primary cancer cells. To this end, we have used primary kidney tumors from 2 patients with c-*met*-mutated PRCC and 3 PRCC patients without c-*met* mutation. The factors KDM4C and BHLHE40 were found to be overexpressed only in the tumors with c-*met* mutation, validating the use of iPSC technology for the analysis of these hereditary cancers. We confirmed that phospho-Met, KDM4C and BHLHE40 were overexpressed in the tumors of two patients with the c-*met*-mutated type 1 PRCC UPN4 and UPN5 (**Fig. 6d, e**). These data demonstrate, for the first time the major interest of the use of iPSC technology to model a hereditary cancer allowing the reproduction of the “cancer in a dish” concept opening the perspective to novel drug testing strategies using simple *in vitro* tests. From this regard, our first results using Sunitinib and cis-Platinum, demonstrate clearly the feasibility of this approach by testing the expression of the KIM-1 protein a known a marker for kidney proximal tubular damage and toxicity in kidney^28^.

**Fig. 6.**
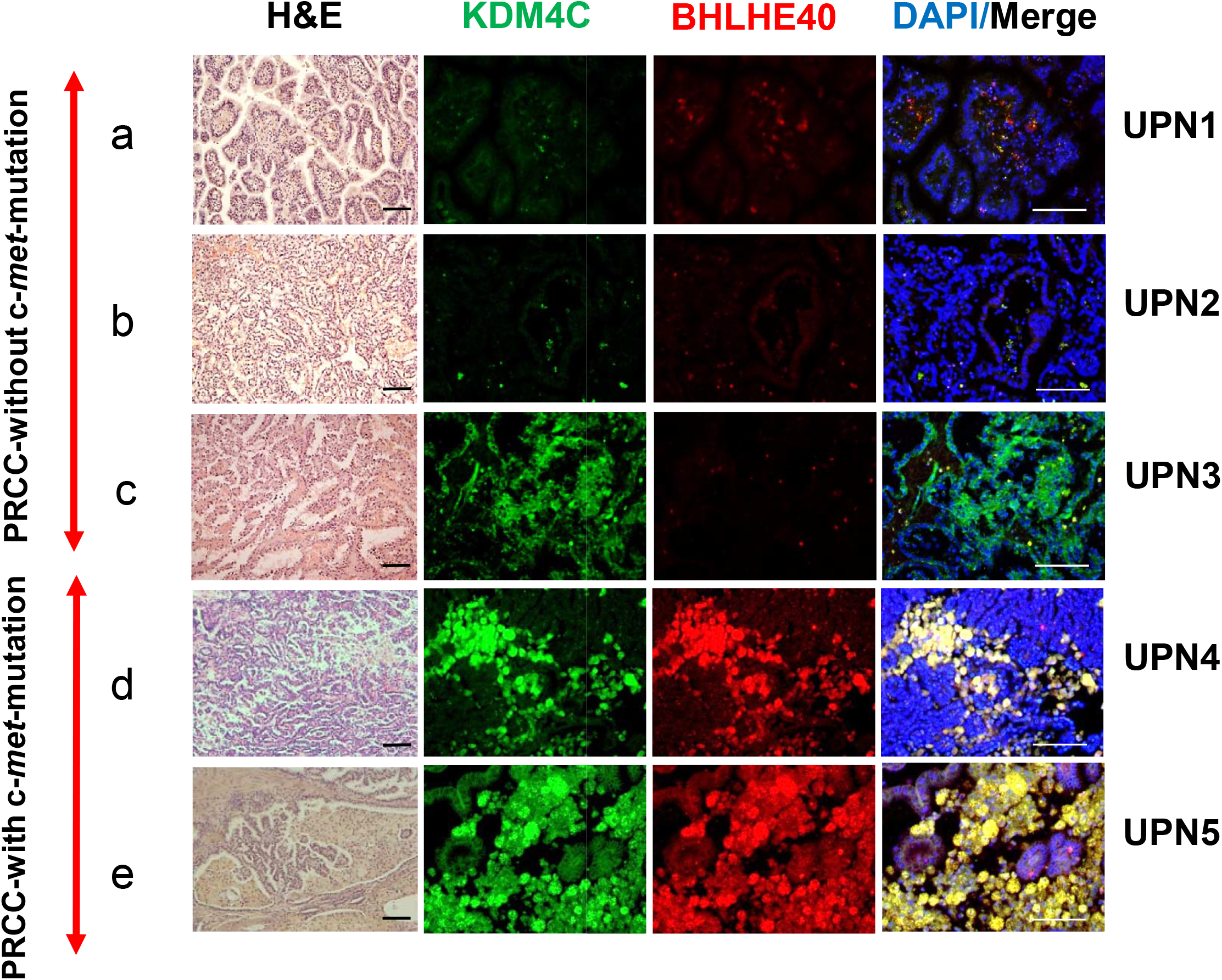
Immunostaining of primary kidney cancer samples from patients without and with c-*Met*-mutated PRCC. Hematoxylin-eosin (H&E) staining and immunostaining for KDM4C, BHLHE40 and DAPI from paraffin-cut primary tumor samples. **a-c** (UPN1, 2, 3) : PRCC without c-*met* mutation. **d,e** (UPN4, 5) : PRCC with c-*met* mutation (see Supplementary Table), Scale bar : 100 μm.

Finally, in order to demonstrate the possible differential behavior *in vivo* of the c-*met*-mutated iPSC derived organoids, we have transplanted these cells as well as control organoids, under the kidney capsule of NSG mice using classical transplantation technology. 4 weeks after transplantation, mice were sacrificed and a necropsy was performed followed by pathological analysis. As can be seen in **Supplementary Fig. 9**, c-*met*-iPSC-derived organoids induced larger tumors as compared to controls and expressed a well known kidney cancer marker TFE3 (transcription factor for immunoglobulin heavy chain enhancer 3)^25^ and Cy7^26^.

In conclusion, we demonstrate in this work for the first time, that c-*met*-mutated hereditary kidney cancer can be modeled in vitro using patient-derived iPSC. This could be achieved by a simple and reproducible 3D-culture system leading to generate kidney organoids without chemically defined induction steps. These organoids could be used for drug toxicity testing. We show that c-*met*-mutated derived-iPSC kidney organoids have the potential to generate the gene profiling similar to that found in primary c-*met*-mutated PRCC, reproducing the expression of several genes known to be expressed in primary cancer cells. Amongst these, KDM4C is a histone demethylase and its overexpression has been shown in several cancers including breast, colon and prostate cancers^29^. In PRCC, CNV of this gene has previously been shown^27^. The pathophysiological role of KDM4C overexpression in c-*met*-mutated PRCC suggest that this pathway can be druggable, as small molecules inhibiting KDM4C are currently been developed^30^. The BHLHE40 (DEC1 / Stra13) is a well-known basic helix loop transcription factor and has been shown to play major roles in cell proliferation, circadian rhythm, tumor progression^31^ and has been shown to be overexpressed in papillary renal cell carcinoma^32^. This work confirms that the discovery of the potential involvement of these pathways is possible by the use of cancer organoids derived from iPSC lines established from hereditary cancers. These findings could open novel perspectives for drug screening as well as future precision medicine strategies in hereditary cancers and could be applied to other hereditary cancers as previously shown in BRCA1-mutated breast^5^ and Li-Fraumeni-syndrome^3^ and RET-mutated^2^ cancers. Finally, in the setting of healthy persons presenting a hereditary cancer-risk mutation, these results could pave the way for the future use of this technology to generate predictive strategies.

## METHODS

### Generation of c-*met*-mutated iPSCs

Bone marrow mononuclear cells (BMNC) were obtained with the informed consent of the patient according to the Declaration of Helsinki and the approval of the Inserm ethical committee. Cell programming was performed using previously reported procedures^2,33^. Briefly, 4 days before reprogramming, BMNC were thawed, cultured and expanded in Myelocult™ medium (Stemcell Technologies) supplemented with 1% penicillin-streptomycin (Life technologies), hSCF 100 ng/mL, hFLT-3100 ng/mL, hIL-3 20 ng/mL, hIL-6 20 ng/mL, and TPO 20 ng/mL (all of them from Peprotech). 2 × 10^5^ BMNC were then transduced overnight with Sendai viruses containing Oct3/4, Sox2, Klf4, and cMyc (CytoTune®-iPS Sendai Reprogramming Kit, Life technologies) each of them at multiplicity of infection (MOI) of 10. The next 2 days, medium was changed daily and cells were resuspended in expansion medium after 5 min centrifugation at 200×g. The third day, cells were recovered by centrifugation and plated on Mitomycin-C-treated mouse embryonic fibroblasts (MEF, CD1 strain) in expansion medium for 2 additional days. At day 6, half of the medium was changed to human pluripotent stem cell medium (hPSC medium) based on DMEM/F12 supplemented with 20% Knock Out Serum Replacer, 1 mM L-glutamine, 1% penicillin/streptomycin, 100 μM 2-mercaptoethanol (all of them from Life technologies) and 12.5 ng/mL basic FGF (Miltenyi Biotech). Then, the medium was changed daily with hPSC medium. At day 26, fully reprogrammed colonies were manually picked and transferred to freshly Mitomycin-C-treated MEF for amplification. iPSC culture was performed according to two different procedures: in the presence of feeders or in feeder-free conditions. Cultures on feeder layers were performed on Mitomycin C-treated MEF as described above with passaging every 7 days using 1 mg/mL collagenase IV in DMEM/F12 (Life technologies) the feeder-free culture was performed on Geltrex^™^ (Life technologies) in essential 8 medium (Life technologies) and 1% penicillin/streptomycin with passaging every 3–4 days using in DPBS (Life technologies) supplemented with 0.5 mM EDTA (Life technologies) and 1.8 mg/L NaCl (Sigma).

### Flow cytometry

Control iPSC or c-*met*-mutated iPSC (c-*met* patient iPSC) colonies were collected using 1 mg/ml collagenaseIV (Life technologies). A single cell suspension was obtained by incubation in Enzyme Free Cell Dissociation Buffer (Life Technologies) followed by mechanical trituration. 100,000 cells were stained in 10 μl PBS supplemented with 1 μl of primary antibodies raised against SSEA4 conjugated to BD Horizon™ V450, SSEA3 conjugated to Phycoerythrin and TRA-1-60 conjugated to Alexa Fluor™ 647 (all of them from BD Biosciences) for 30 minutes at 4°C. Cells were then washed with PBS and analyzed using a MACSQuant flow cytometer (MiltenyiBiotec).

### Teratoma assays

Animal experimentation was performed according to French regulations. Protocols were approved by the Ethical Committee for Animal Experimentation (CEEA n°26) under the agreement number 2015-012-534. Experiments were performed using female mice aged 6 to 7 weeks old. Control iPSC or c-*met*-mutated iPSC were collected by collagenase IV treatment (Life technologies). 2 × 10^6 cells were resuspended in 150 μl of Geltrex^™^:DMEM/F12 (1:1) and were subcutaneously injected into the rear leg of NSG mice (NOD.Cg-Prkdcscid Il2rgtm1Wjl/SzJ). After 8 or 12 weeks, teratomas were isolated and processed for histological analysis.

### Cell culture and generation of cell aggregates

Control and c-*met*-mutated iPSC were maintained on Geltrex (Stem Cell Technologies, Inc) coated flat culture dish in E8 media (Stem Cell Technologies, Inc) contained DMEM/F12, L-ascorbic acid-2-phosphate magnesium (64 mg/l), sodium selenium (14 μg/l), FGF2 (100 μg/l), insulin (19.4 mg/l), NaHCO3 (543 mg/l) and transferrin (10.7 mg/l), TGFβ1(2 μg/l) or NODAL (100 μg/l). Osmolarity of all media was adjusted to 340 mOsm at pH7.4. All the media were stored at 4°C, and were used within 2 weeks of production. Colonies were manually harvested at 60-80% confluence. Cells were then collected and dissociated into single cells using EDTA. Cells (1×10^6 or 1×10^5/well) were put onto 24 or 96-well ultralow attachment plates (Corning, Inc.) to allow them to form aggregates in suspension in a CO2 incubator at 37°C, in 5% CO2. Cell aggregates were cultured in E8 medium (Stem Cell Technologies) with daily medium change for 1-7 days.

### Generation of kidney organoids

Control or c-*met*-mutated iPSC-aggregates (iPSC-A) were plated on a Geltrex (Stem Cell Technologies, Inc.) in 24 or 96-well plates or 8-well culture chambers. The aggregates were cultured in E8 medium (STEMCELL Technologies, Inc.) with daily medium change for 6-7 days. Photographies were taken using a NIKON microscope.

### Immunofluorescence

Kidney organoids generated in vitro or tumors arising in NSG mice were embedded in paraffin. Paraffin sections were de-paraffinized and permeabilized with 0.2% Triton X-100 (Sigma) in PBS and blocked in 10% serum. Primary antibodies were diluted in PBS containing 10% serum at the following concentrations: PODXL; 1:200 (Abcam), LTL; 1:200 (Clinisciences), Cytokeratin 7; 1:200 (Abcam), VHL; 1:200 (Santa Cruz), phospho-Met; 1:200 (Ozyme), KDM4C; 1:200 (Abcam), TFE3; 1:200 (Abcam) and washed three times in PBS. The sections were incubated with secondary antibodies in antibody dilution buffer for 1 hour, then washed three times in PBS. Nuclei were labeled with DAPI mounting medium. Visualization and capture were realized with a Zeiss confocal microscope and Volocity software or NIKON microscope.

### Whole-mount immunohistochemistry of 3D kidney organoids

iPSC-A or kidney organoids cultured on 96-well culture dishes or 8-well culture chambers were washed with phosphate-buffered saline (PBS), fixed with 4% paraformaldehyde in PBS for 120 min, permeabilized with 0.2% Triton X-100 (Sigma) in PBS and blocked in 10% serum. Primary antibodies were diluted in PBS 10% serum at the following concentrations: Brachyury; 1:200, PODXL; 1:200 (Abcam), LTL; 1:200 (Clinisciences), Six2; 1:200 (Euromedex), TFE3; 1:200 (Abcam), Cytokeratin 7; 1:200 (Abcam), Nephrin; 1:200 (Abcam), KIM-1; 1:200 (Biotechne), Phalloidin; 1:200, phospho-Met; 1:200 (Ozyme), KDM4C; 1:200 (Abcam) and then washed in PBS. The samples were incubated with secondary antibodies in antibody dilution buffer, then washed in PBS. Nuclei were labeled with DAPI mounting medium. Visualization and capture were realized with a Zeiss confocal microscope and Volocity software or NIKON microscope.

### Transmission electron microscopy (TEM)

Kidney organoids were gently centrifuged and pelleted before the TEM process as follows. The cells were fixed in 2.5% glutaraldehyde in phosphate-buffered saline (PBS) for 1h at 4°C, washed in PBS, and fixed in 1% osmium tetroxide in PBS for 1h. They were dehydrated in ascending series of graded ethyl alcohols, then in acetone. Each sample was infiltrated with the resin before being embedded in epoxy resin and polymerized for 72h. Semi-thin sections of about 0.5 to 1 μm were obtained and colored with Toluidine blue before being examined via a light microscope with an associated digital camera, hooked to a computer for image processing and editing (Leica DC 300). Ultra thin sections of about 60/90 nm were contrasted with heavy metals (uranyl acetate and lead citrate) and were examined using a Jeol 1010 transmission electron microscope at an accelerated voltage of 80kV. Images were photographed on digital images Gatan Digital Micrograph : brure Erlangen 500w : camera and edited by Microsoft Power Point.

### Functional analysis for proximal tubules of kidney organoids

For dextran uptake assay, kidney organoids at day+12 were cultured with 20 μg/ml of 10,000MW dextran Alexa 488-conjugated (D-22910, Life Technologies) for 48 h. Organoids were 4% PFA fixed, then washed three times in PBS. Nuclei were labeled with DAPI mounting medium. Visualization and capture were realized with a NIKON microscope.

### Toxicity tests using c-*met*-mutated iPSC-derived kidney organoids

Control or c-*met*-mutated iPSC-derived kidney organoids cultured in 96-well plates were treated with cisplatin (5000 μg/ml) for 24h or with sunitinib at low (200 μg/ml) or high (5000 μg/ml) concentration for 96h. They were then fixed, and processed for immunofluorescence with KIM-1 antibodies (Biogen, 1:100 dilution).

### Kidney intra-capsule transplantation experiments

Kidney organoids cultured for 12–14 days were transplanted beneath the renal capsule of 30–32 week-old male NSG mice (NOD.Cg-Prkdcscid Il2rgtm1Wjl/SzJ). 4 weeks after transplantation, mice were euthanized, kidneys were removed and processed for histology and immunofluorescence analysis.

### Kidney histology and immunofluorescence analyses

After euthanasia, the kidneys containing the transplanted organoids were fixed in paraformaldehyde and embedded in paraffin. 4 μm sections were deparaffinized and then permeabilized with 0.2% Triton X-100 (Sigma) in PBS and blocked 10% serum. Primary antibodies were diluted in PBS 10% serum at the following concentrations: PODXL; 1:200 (Abcam), LTL; 1:200 (Clinisciences), Cyto-keratin 7; 1:200 (Abcam), TFE3; 1:200 (Abcam), Nephrin; 1:200 (Abcam) then washed three times in PBS. The sections were incubated with secondary antibodies in antibody dilution buffer for 1 hour, then washed three times in PBS. Nuclei were labeled with DAPI mounting medium. Visualization and capture were realized with a Zeiss confocal microscope and Volocity software or NIKON microscope.

### Immunochemistry analysis of primary kidney tumors

Two patients with c-*met*-mutated PRCC (Tumor Samples UPN4, UPN5) and primary PRCC tumors from three patients without c-*met* mutation (Tumor samples UPN1, UPN2 and UPN3) kidney tissue were embedded in paraffin. Paraffin sections were deparaffinized and then permeabilized with 0.2% Triton X-100 (Sigma) in PBS and blocked 10% serum. Primary antibodies were diluted in PBS 10% serum at the following concentrations: PODXL; 1:200 (Abcam), phospho-Met; 1:200 (Ozyme), KDM4C; 1:200 (Abcam), TFE3; 1:200 (Abcam) and then washed three times in PBS. The sections were incubated with secondary antibodies in antibody dilution buffer for 1 hour, then washed three times in PBS. Nuclei were labeled with DAPI mounting medium. Visualization and capture were realized with a NIKON microscope

### Public datasets

RNAseq and Genomic experiments from dataset of papillary renal cell carcinoma from TCGA consortium^14^ were downloaded from Cbioportal database access center^34^. This analyzed dataset comprised 291 samples of PRCC tumors quantified by RNAseq at level V2 Z-scores – 21 patients were found mutated for c-*met* in this cohort (Supplemental Table 4).

### Transcriptome analyses of iPSC-A

Total RNA from c-*met*-mutated iPSC (monolayer culture) and c-*met*-mutated iPSC-A (iPSC aggregates) was extracted isolation RNA with the PureLink RNA Mini Kit (Life technology) by following manufacturer instructions. Quantification of RNA was performed with Nanodrop technology and RNA integrity was controlled with Agilent Bioanalyzer (Agilent technologies). Microarray probes were synthetized and labelled by linear amplification kit as Affymetrix manufacturer instructions to be hybridized to Human Clariom S microarray compatible with Affymetrix station. Resulting scanned files (CEL files) were normalized with RMA method with Expression console version 1.4.6 (Affy*met*rix)^35^.

### Bioinformatics

Bioinformatics analysis was performed in R software environment version 3.0.2. Ranking product analysis was performed on transcriptome controlized matrix with implementation of one hundred of permutations^36^ supervised by class description defined by culture conditions of the iPSC: c-*met*-mutated iPSC (monolayer culture) versus c-*met*-mutated iPSC-A. Expression heatmap was performed with made 4 bioconductor library by using Euclidean distances and Ward classification method^37^. Analysis of PRCC RNAseq dataset was performed by machine learning with library pamr^38^. Protein-protein interaction network was built with Network Analy-stapplication^39^ on innateDB interaction database^40^. Functional inference on interaction network was enriched with Kyoto Encyclopedia of Genes and Genomes (KEGG) database^41^.

### Transcriptome analyses of iPSC-derived kidney organoids

Microarray Clariom S human was done on process total RNA from kidney organoids derived samples (c-*met*-mutated iPSC and control iPSC) in duplicates. Expression matrix was built with CEL files generated on Affymetrix Station and normalized by RMA method with Expression console software (Affymetrix) version 1.4.1.46^35^. Gene set enrichment analysis was performed between conditions with MsigDb database version 6.0^42^.

## AUTHOR CONTRIBUTIONS

JWH and AGT conceived, designed, analyzed data and wrote the manuscript. JJP followed the patients in the Urology Department. SR performed genetic c-*met*-mutation-detection experiments. OF generated iPSCs. JWH performed all iPSC-derived organoid experiments. JLD performed TEM experiments. CD analyzed bioinformatics data. AF performed transplantation experiments under the NSG renal capsule. SF and VW performed pathology analyses. AGT, ABG, FG conceived, designed, analyzed data and supervised the project. JWH, AF and AGT wrote the manuscript.

## SUPPLEMENTARY FIGURES

**Supplementary Fig. 1.**
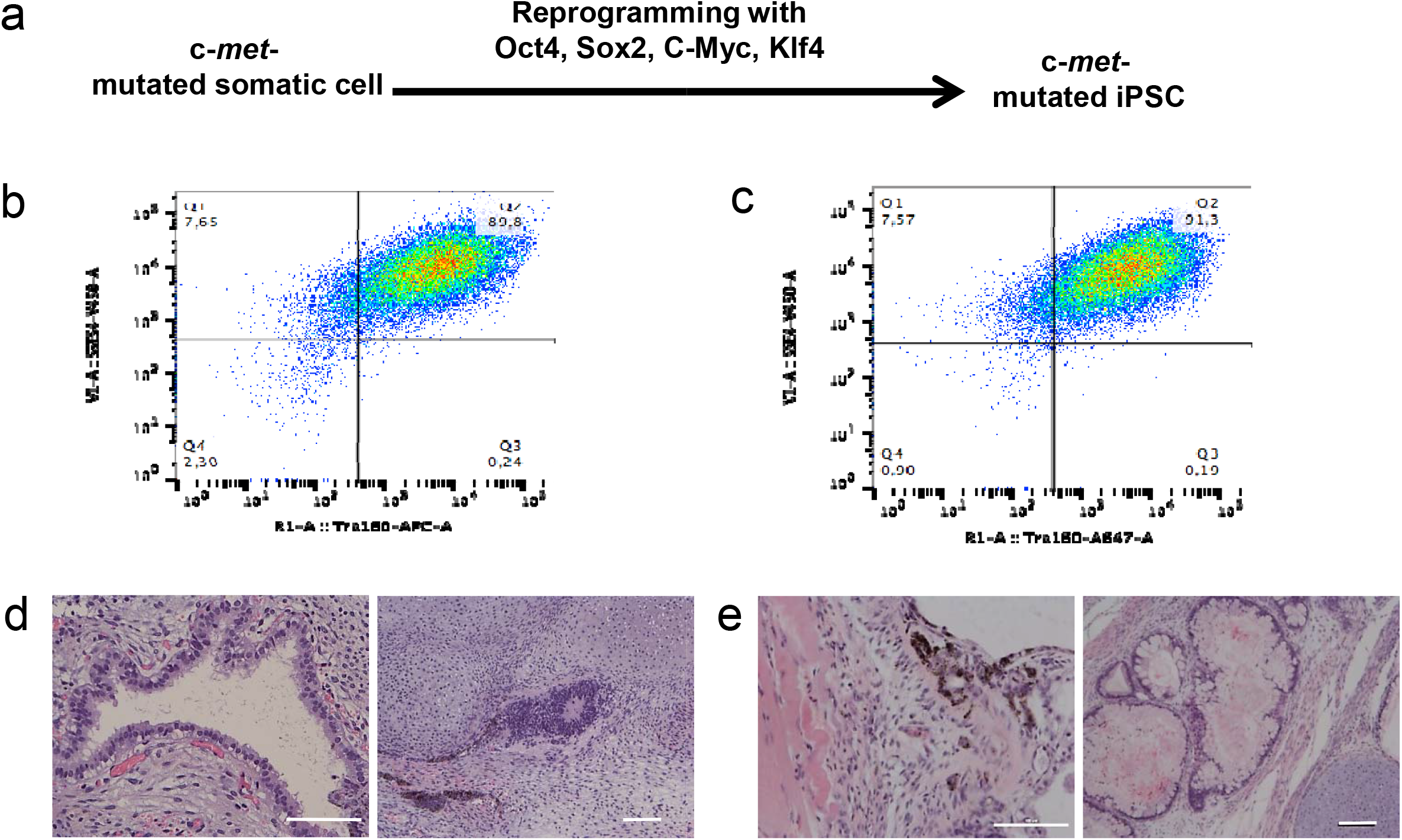
Generation and characterization of c-*met*-mutated iPSC using immunophenotyping and teratoma assays. **a,** Protocol used for the generation of c-*met* iPSC using Sendaï-virus mediated - mediated pluripotent gene transfer. **b,c** FACS analysis of control (left panel) or c-*met*-mutated iPSC (right panel) using pluripotency markers Tra-1-60 and SSEA4. **d,e** H&E staining section of control (left panel) or c-*met*-mutated iPSC (right panel) showing derivatives of all three germ layers. Scale bar : 100 μm.

**Supplementary Fig. 2.**
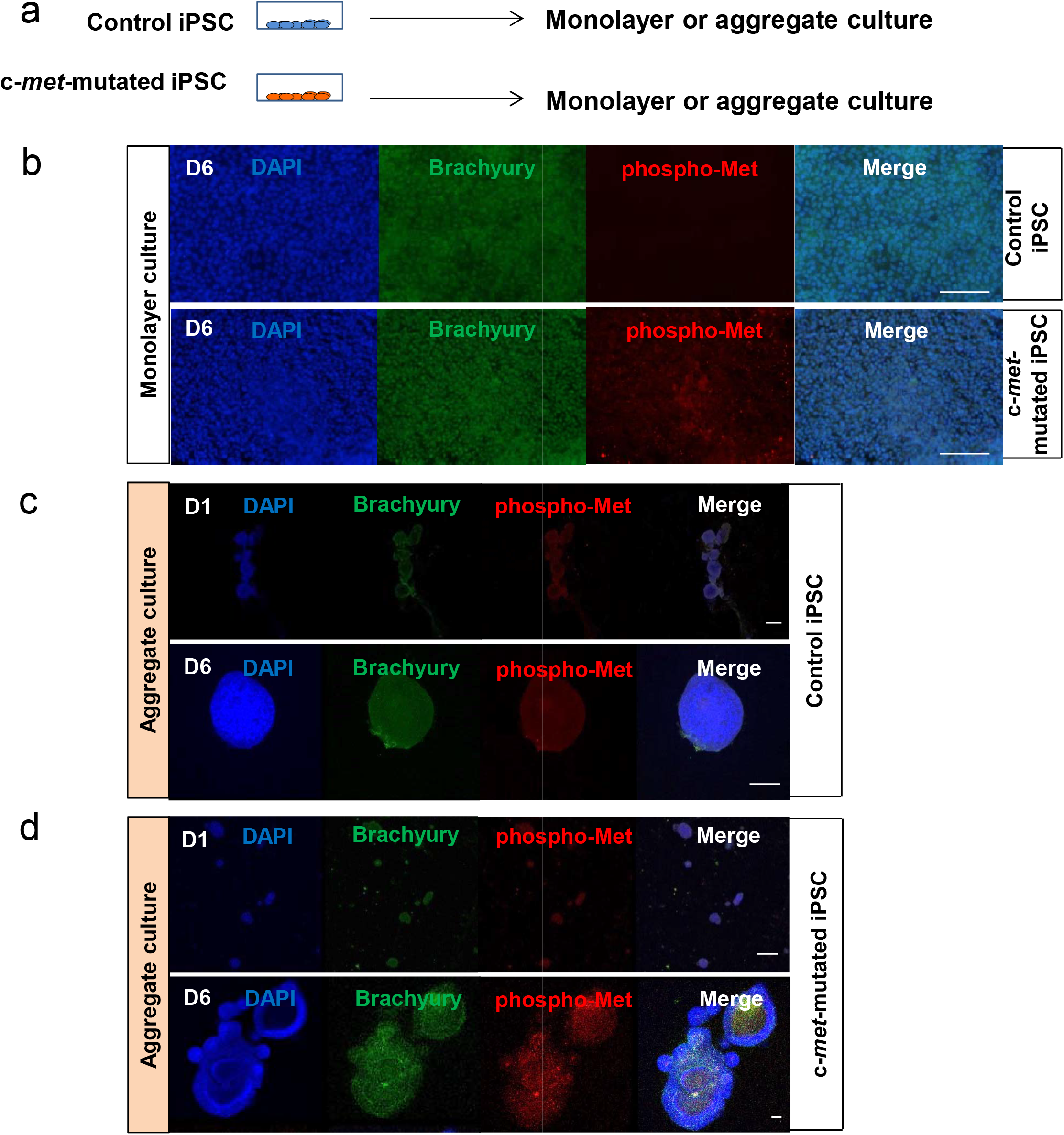
Immunochemistry analyses of c-*met*-mutated iPSC aggregates and evaluation of phospho-MET expression. **a,** Schematic representation of control and c-*met*-mutated iPSC grown in monolayer cultures with and without low attachment conditions generating aggregates. **b,** Immunochemistry for Brachyury, phospho-Met and DAPI on day 6 of control iPSC or c-*met*-mutated iPSC monolayer culture. A faint phospho-MET expression is detected in c-*met*-mutated aggregates. **c,d,** Immunocytochemistry analyses for Brachyury and phospho-Met expression on day 1 and 6 of control **(c)** or c-*met*-mutated iPSC aggregates revealing similar levels of phospho-MET expression in both conditions. Scale bar : 100 μm.

**Supplementary Fig. 2.**
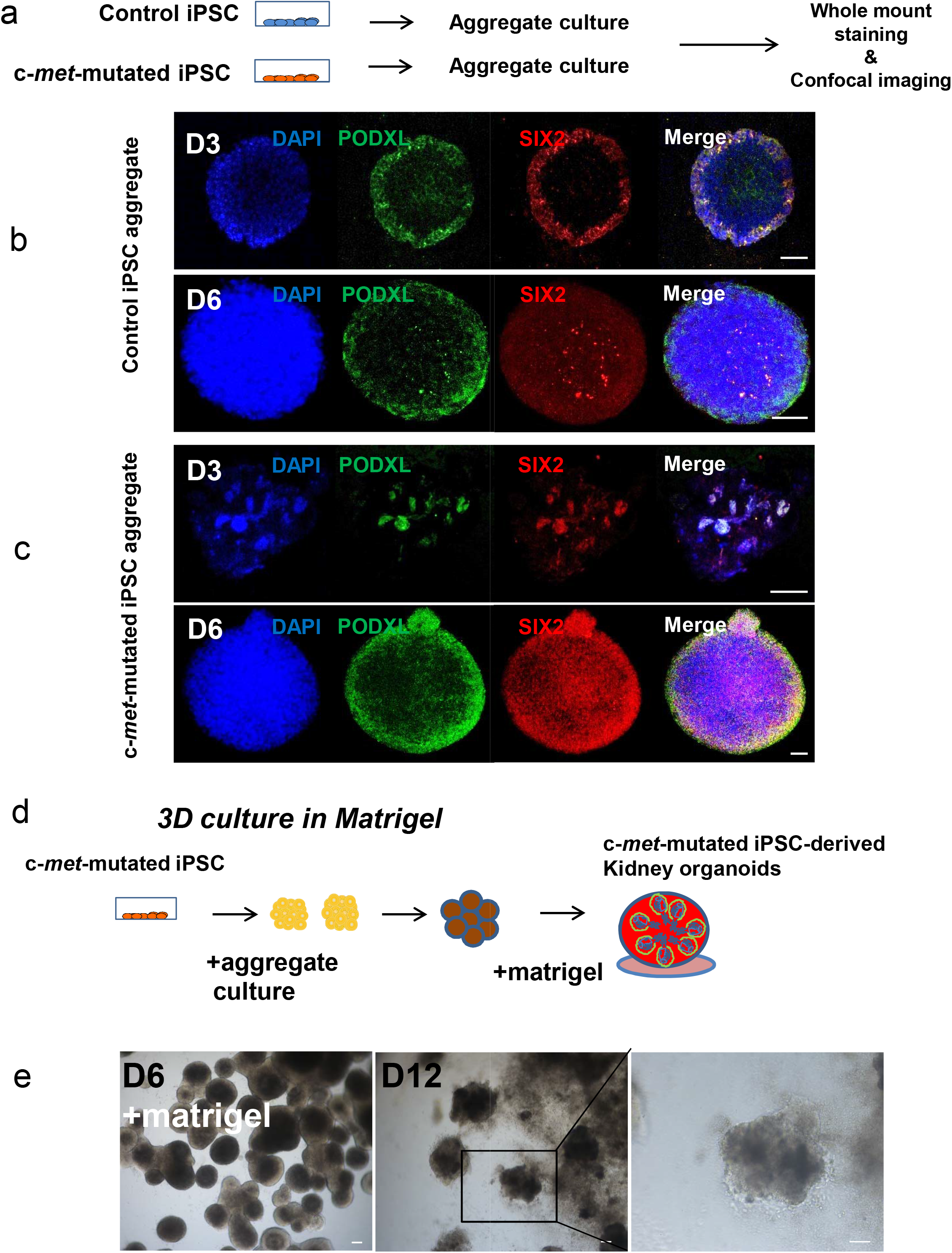
Appearance of kidney differentiation markers in aggregate cultures to optimize kidney organoid generation in 3D cultures. **a,** Schematic representation of the experiments designed to generate control iPSC and c-*met*-mutated iPSC in aggregate cultures performed in low attachment plates. **b,c** Whole-mount staining for PODXL, Six2 and DAPI of control iPSC aggregates on day 3 and day 6 showing the appearance of kidney markets at day+3, with increased expression at day+6 in both control and c-*met*-mutated cells. Scale bar : 50 μm. **d** Schema of the protocol used for of 3D cultures in Matrigel, based on the results obtained in aggregate cultures withy optimal kidney differentiation at day+6. **e:** 3D cultures at day+6 in the presence of Matrigel allowed the generation structures in which kidney differentiation markers were found (See Figure S4) Scale bar : 100 μm.

**Supplementary Fig. 4.**
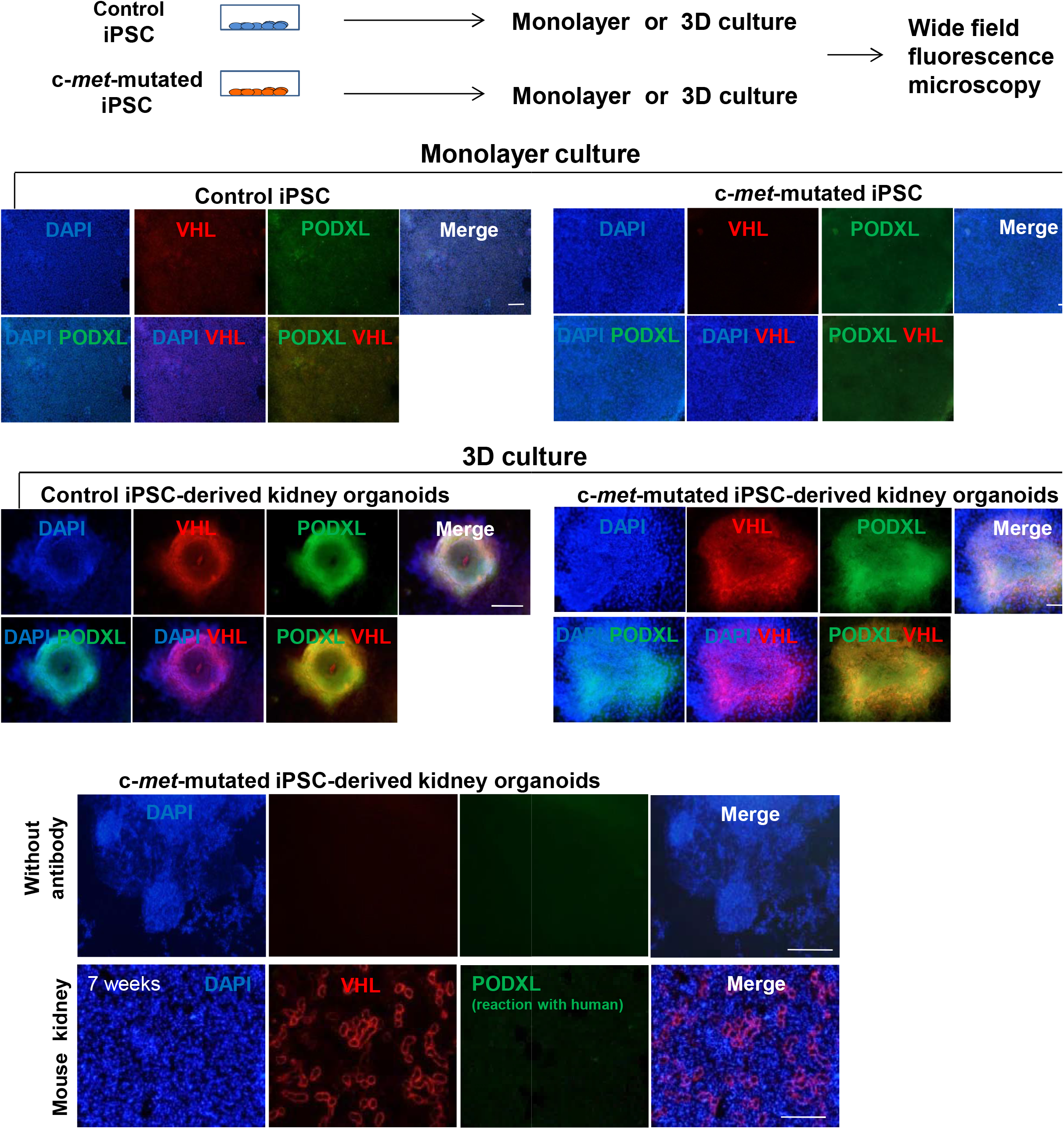
Self-organizing kidney organoids are detected only in 3D cultures. **a,** Schematic representation of experiments designed performed in control iPSC and c-*met*-mutated iPSC monolayer or 3D culture. **b,** Immunocytochemistry for PODXL, VHL and DAPI of iPSC in monolayer culture on day 6. The use of PODXL antibodies does not allow the detection of kidney differentiation. **c,** Whole-mount staining of organoids in 3D cultures followed by staining with PODXL, VHL allows detection of kidney differentiation markets at day+12. **D: Control experiment showing** whole-mount staining without 1^st^ antibody of c-*met*-mutated iPSC-derived kidney organoids 3D culture on day 12. **e,** Immunohistochemistry for PODXL, VHL and DAPI of mouse kidney. Scale bar : 100 μm.

**Supplementary Fig. 5.**
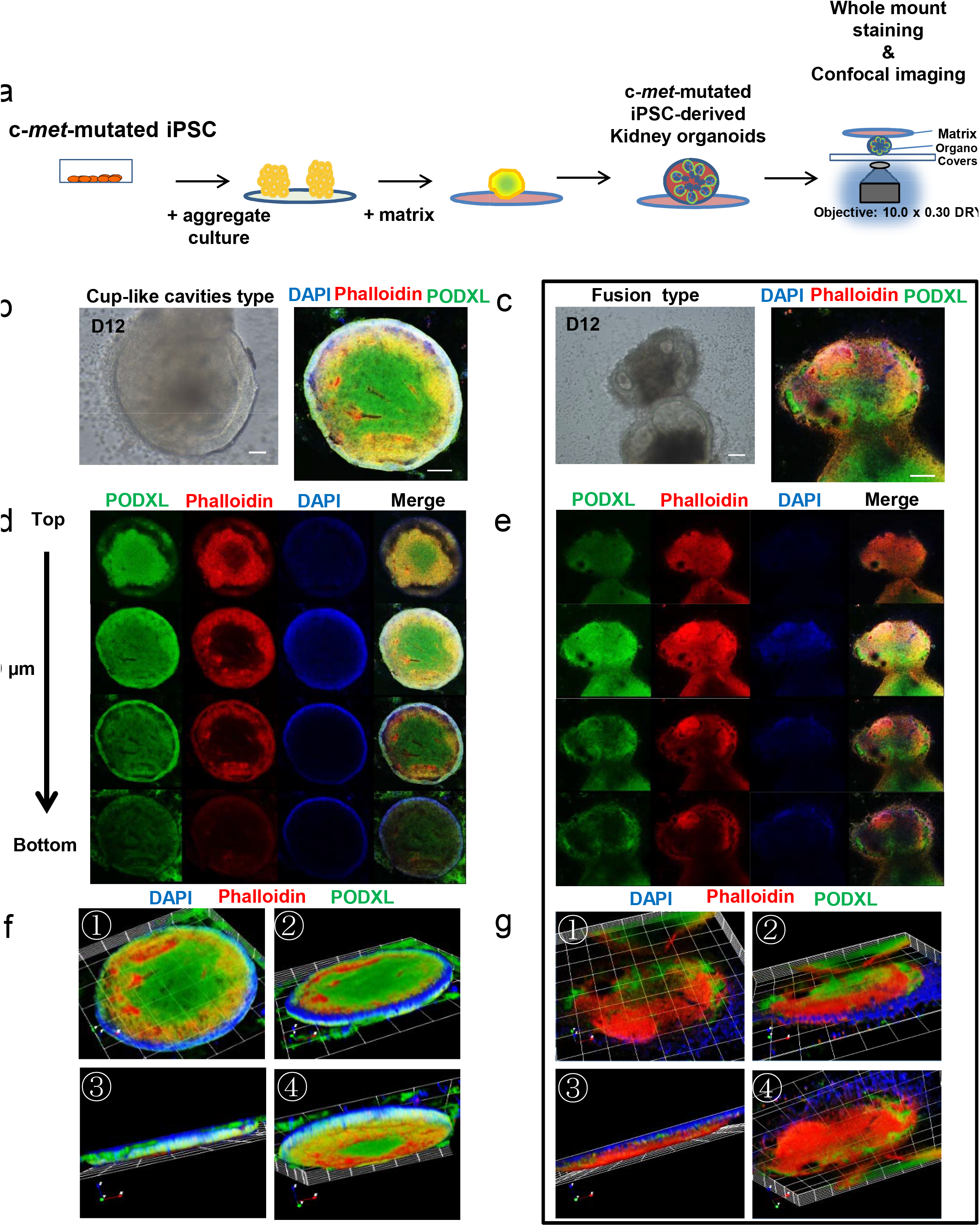
Self-organizing iPSC-derived kidney organoids. **a,** Schematic representation of the experiments designed to generate c-*met*-mutated iPSC in aggregate culture. **b,c,** Optical image of c-*met*-mutated iPSC-derived kidney organoids and confocal image of whole-mount staining for PODXL, phalloidin and DAPI, Scale bar : 100 μm. **d,e,** Confocal image of cup-like cavities and fusion type. **f,g,** 3D rotation image of cup-like cavities and fusion type.

**Supplementary Fig. 6.**
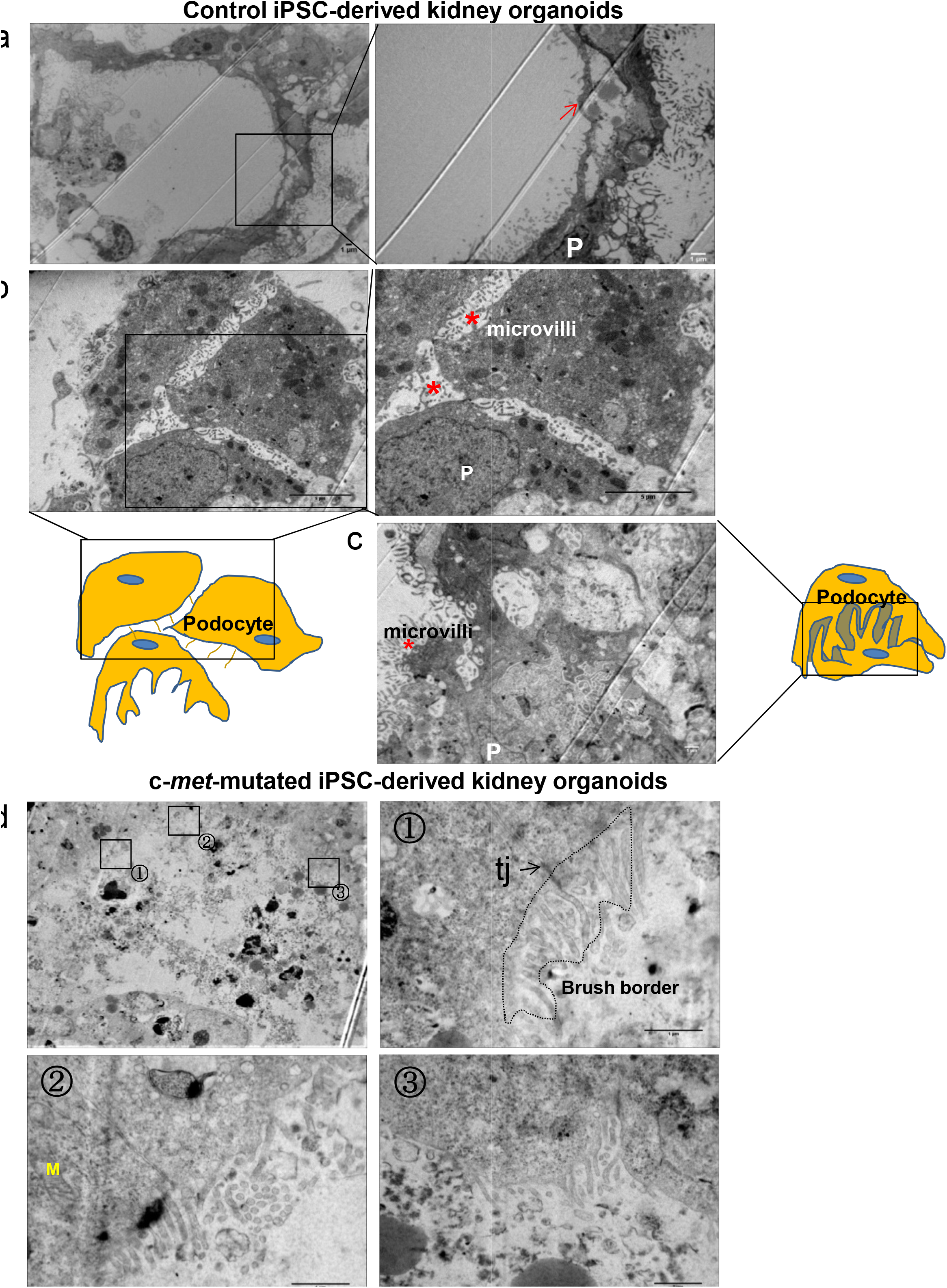
Electron microscopy of iPSC-derived kidney organoids. **a,b,c,** Representative electron microscopy images glomerulus region of control iPSC-derived kidney organoids, podocyte-like cells (P), glomerular basement membrane (arrow), microvilli (*). **d,** Representative electron microscopy images tubule region of c-*met*-mutated iPSC-derived kidney organoids, tight junctions (tj), mitochondria (M).

**Supplementary Fig. 7.**
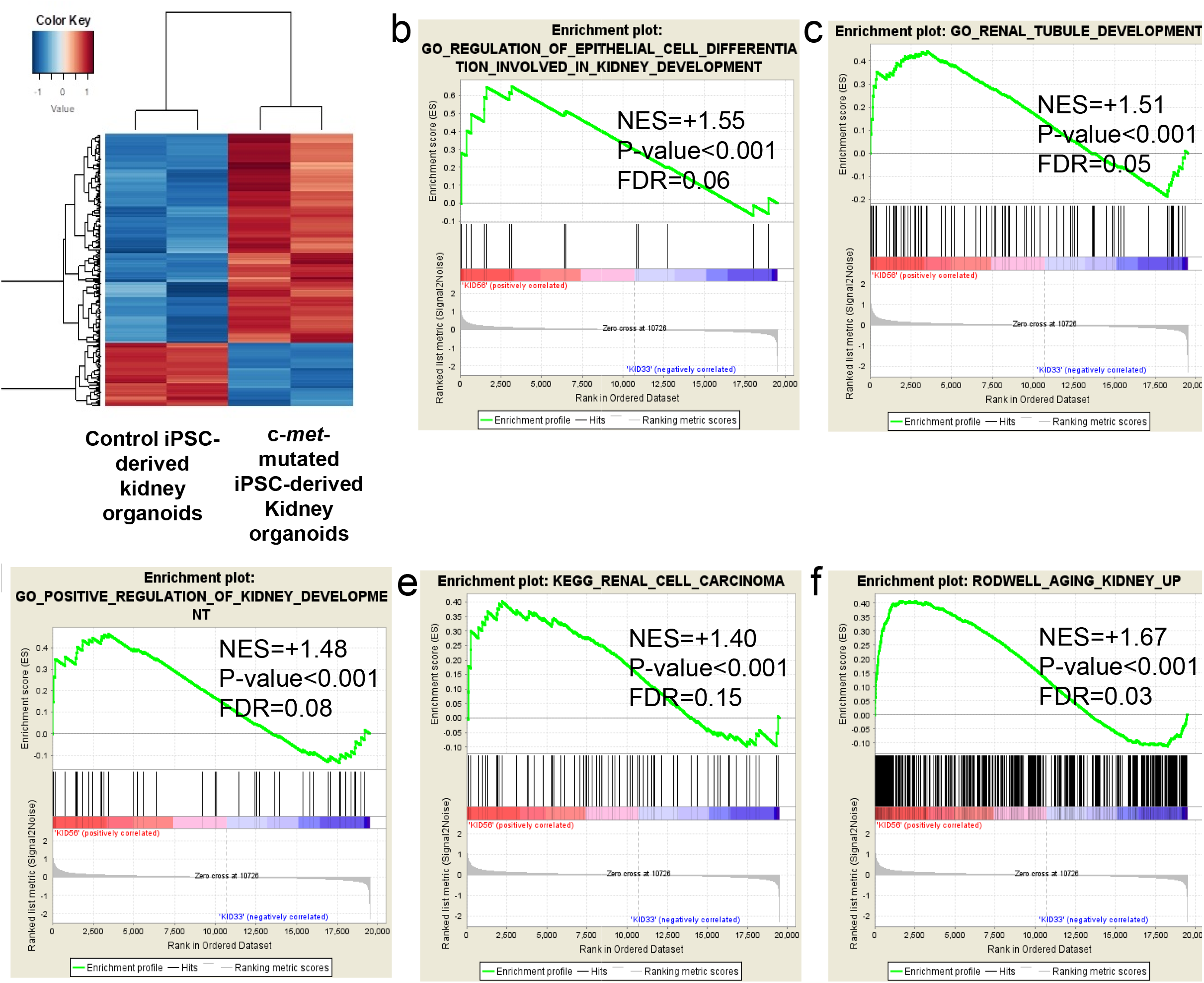
Functional enrichment of differentialy expressed genes between control versus c-*met*-mutated iPSC-derived kidney organoids. **a,b,c,** Kidney development gene sets enriched in c-*met*-mutated iPSC-derived kidney organoids versus control iPSC-derived kidney organoids, NES: normalized enriched score, FDR: False Discovery Rate. **d,e,** Gene sets for kidney pathophysiological enriched in c-*met*-mutated iPSC-derived kidney organoids. versus control iPSC-derived kidney organoids, NES: normalized enriched score, FDR: False Discovery Rate.

**Supplementary Fig. 8.**
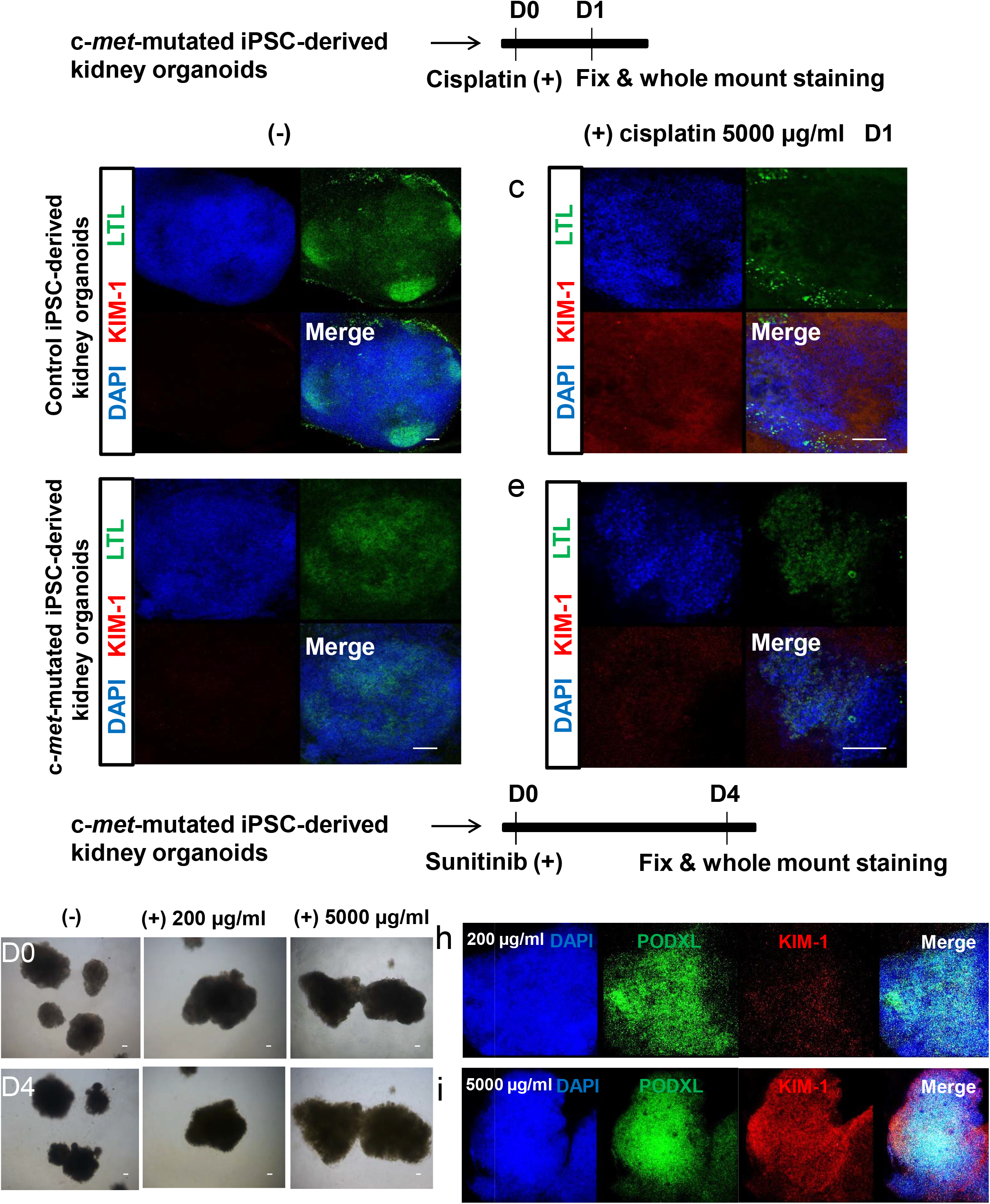
Drug toxicity experiments. **a,** Schematic representation of cisplatin toxicity experiments in c-*met*-mutated iPSC-derived kidney organoids. **b,** Representative whole-mount staining for LTL, KIM-1 and DAPI in control iPSC-derived kidney organoids treated with cisplatin (5000 μg/ml) (right panels). **c,** Representative whole-mount staining for LTL, KIM-1 and DAPI in c-*met*-mutated iPSC-derived kidney organoids treated with cisplatin (5000 μg/ml) (right panels). **d,** Schematic of drug sunitinib toxicity test process of c-*met*-mutated iPSC-derived kidney organoids. **e,** Photograph of c-*met*-mutated iPSC-derived kidney organoids in 96 well plate. **f,** Representative whole-mount staining for PDOXL, KIM-1 and DAPI in c-*met*-mutated iPSC-derived kidney organoids treated with sunitinib (200 μg/ml). **g,** Representative whole-mount staining for PODXL, KIM-1 and DAPI in c-*met*-mutated iPSC-derived kidney organoids treated with sunitinib (5000 μg/ml), Scale bar : 100 μm.

**Supplementary Fig. 9.**
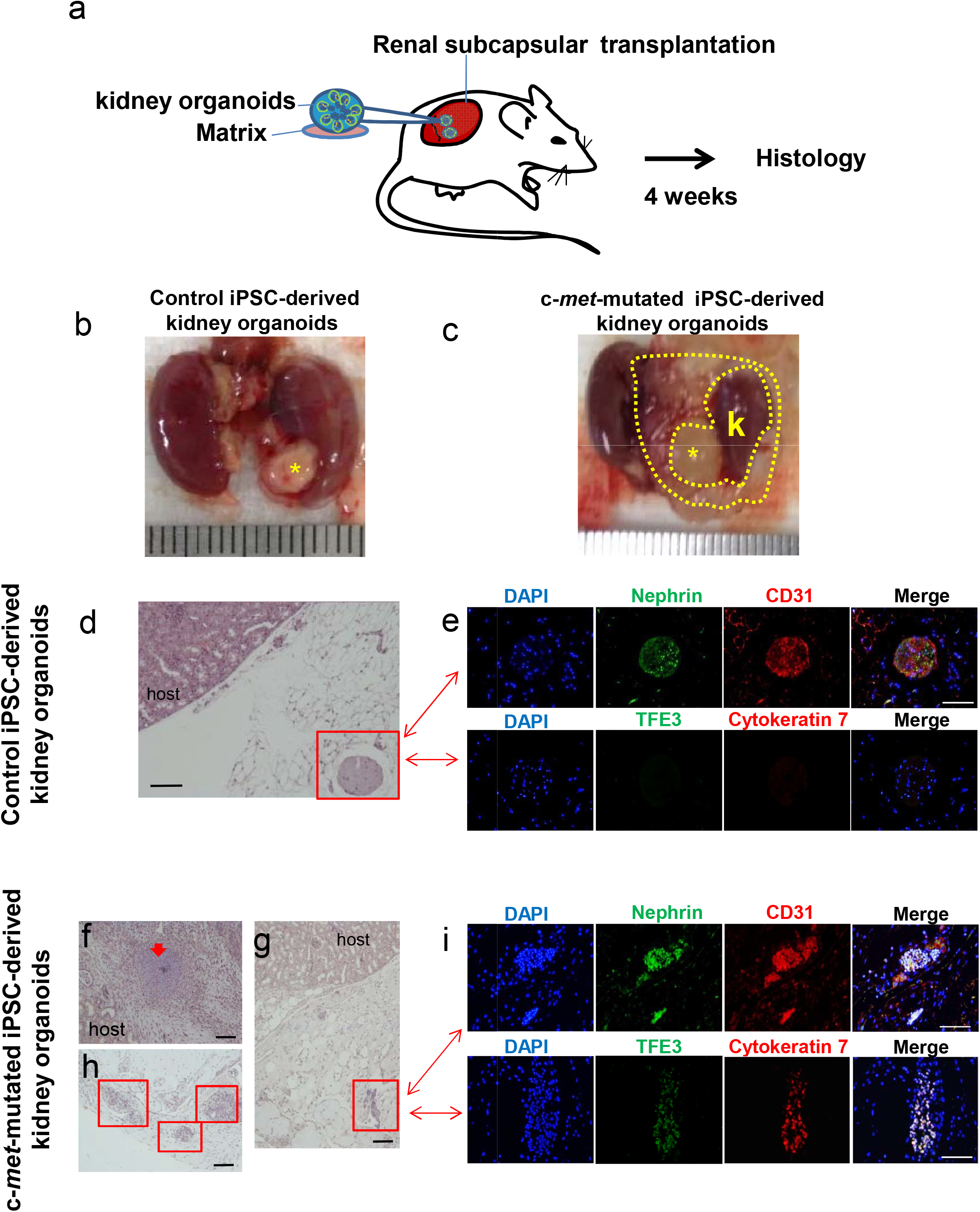
Analysis of tumors generated by transplantation of kidney organoids. **a.** Experimental protocol used. **b,c:** Macroscopic aspect of kidney NSG mouse 1 month after transplantation of control (*) or c-*met*-mutated iPSC-derived kidney organoids** **d,e** Pathological analysis of tumors genereated by transplantation of control kidney organoids at 1 month showing the presence of normal organoid-like structures expressing kidney differentiation markers. **f,h,g,i** Pathology of c-*met*-mutated tumors after transplnation of c-*met*-mutated kidney organoids at 4 weeks, showing disorganized structures with expression of kidney cancer markers TFE3 and Cy7. Scale bar : 100 μm.

## SUPPLEMENTARY TABLES

**Supplementary Table 1.**

List of genes differentially expressed between c-*met*-mutated iPSC-derived kidney organoids and control iPSC-derived kidney organoids.

**Supplementary Table 2**

List of genes with differential expression between c-*met*-mutated iPSC aggregates versus c-*met*-mutated iPSC: red rows described genes which were found up regulated in c-*met*-mutated iPSCs aggregates as compared to c-*met*-mutated iPSC and green rows described genes which were found down regulated in c-*met*-mutated iPSCs aggregate as compared to c-*met*-mutated iPSC.

**Supplementary Table 3**

List of genes whose expression identified during meta-analysis in both iPSCs transcriptome study and RNA-sequencing performed on PRCC patients with c-*met* mutation. In red : positive fold change; In green : negative fold change.

**Supplementary Table 4.**

List of patients with c-*met* mutations in the TCGA cohort of papillary renal-cell cancer.

## REFERENCES

1. Cragun, D. & Pal, T. Identification, Evaluation, and Treatment of Patients with Hereditary Cancer Risk within the United States. ISRN Oncol. 2013, 260847 (2013).

2. Hadoux, J. et al. Generation of an induced pluripotent stem cell line from a patient with hereditary multiple endocrine neoplasia 2A (MEN2A) syndrome with RET mutation. Stem Cell Res. 17, 154–157 (2016).

3. Lee, D.-F. et al. Modeling familial cancer with induced pluripotent stem cells. Cell 161, 240–254 (2015).

4. Griscelli, F. et al. Generation of induced pluripotent stem cell (iPSC) line from a patient with triple negative breast cancer with hereditary exon 17 deletion of BRCA1 gene. Stem Cell Res. 24, 135–138 (2017).

5. Soyombo, A. A. et al. Analysis of Induced Pluripotent Stem Cells from a BRCA1 Mutant Family. Stem Cell Rep. 1, 336–349 (2013).

6. Kim, J. et al. An iPSC line from human pancreatic ductal adenocarcinoma undergoes early to invasive stages of pancreatic cancer progression. Cell Rep. 3, 2088–2099 (2013).

7. Huang, L. et al. Ductal pancreatic cancer modeling and drug screening using human pluripotent stem cell- and patient-derived tumor organoids. Nat. Med. 21, 1364–1371 (2015).

8. Chartier, S. et al. Biphasic Squamoid Alveolar Renal Cell Carcinoma: 2 Cases in a Family Supporting a Continuous Spectrum With Papillary Type I Renal Cell Carcinoma. Am. J. Surg. Pathol. 41, 1011 (2017).

9. Self, M. et al. Six2 is required for suppression of nephrogenesis and progenitor renewal in the developing kidney. EMBO J. 25, 5214–5228 (2006).

10. Kobayashi, A. et al. Six2 defines and regulates a multipotent self-renewing nephron progenitor population throughout mammalian kidney development. Cell Stem Cell 3, 169–181 (2008).

11. Delahunt, B. & Eble, J. N. Papillary renal cell carcinoma: a clinicopathologic and immunohistochemical study of 105 tumors. Mod. Pathol. Off. J. U. S. Can. Acad. Pathol. Inc 10, 537–544 (1997).

12. Gremel, G. et al. A systematic search strategy identifies cubilin as independent prognostic marker for renal cell carcinoma. BMC Cancer 17, (2017).

13. Tsuda, M. et al. TFE3 Fusions Activate MET Signaling by Transcriptional Up-regulation, Defining Another Class of Tumors as Candidates for Therapeutic MET Inhibition. Cancer Res. 67, 919–929 (2007).

14. Cancer Genome Atlas Research Network et al. Comprehensive Molecular Characterization of Papillary Renal-Cell Carcinoma. N. Engl. J. Med. 374, 135–145 (2016).

15. Lubensky, I. A. et al. Hereditary and Sporadic Papillary Renal Carcinomas with c-met Mutations Share a Distinct Morphological Phenotype. Am. J. Pathol. 155, 517–526 (1999).

16. Siegel, R. L., Miller, K. D. & Jemal, A. Cancer statistics, 2016. CA. Cancer J. Clin. 66, 7–30 (2016).

17. Lam, A. Q. et al. Rapid and efficient differentiation of human pluripotent stem cells into intermediate mesoderm that forms tubules expressing kidney proximal tubular markers. J. Am. Soc. Nephrol. JASN 25, 1211–1225 (2014).

18. Takasato, M. et al. Kidney organoids from human iPS cells contain multiple lineages and model human nephrogenesis. Nature 526, 564–568 (2015).

19. Morizane, R. et al. Nephron organoids derived from human pluripotent stem cells model kidney development and injury. Nat. Biotechnol. 33, 1193–1200 (2015).

20. Dutta, D., Heo, I. & Clevers, H. Disease Modeling in Stem Cell-Derived 3D Organoid Systems. Trends Mol. Med. 23, 393–410 (2017).

21. Davidowitz, E. J., Schoenfeld, A. R. & Burk, R. D. VHL Induces Renal Cell Differentiation and Growth Arrest through Integration of Cell-Cell and Cell-Extracellular Matrix Signaling. Mol. Cell. Biol. 21, 865–874 (2001).

22. Luckett, W. P. The development of primordial and definitive amniotic cavities in early rhesus monkey and human embryos. Am. J. Anat. 144, 149–167 (1975).

23. Takabatake, Y. et al. The CXCL12 (SDF-1)/CXCR4 axis is essential for the development of renal vasculature. J. Am. Soc. Nephrol. JASN 20, 1714–1723 (2009).

24. Yu, W. et al. Clinicopathological, genetic, ultrastructural characterizations and prognostic factors of papillary renal cell carcinoma: New diagnostic and prognostic information. Acta Histochem. 115, 452–459 (2013).

25. Chevarie-Davis, M. et al. The morphologic and immunohistochemical spectrum of papillary renal cell carcinoma: study including 132 cases with pure type 1 and type 2 morphology as well as tumors with overlapping features. Am. J. Surg. Pathol. 38, 887–894 (2014).

26. Garcia, J. & Lizcano, F. KDM4C Activity Modulates Cell Proliferation and Chromosome Segregation in Triple-Negative Breast Cancer. Breast Cancer Basic Clin. Res. 10, 169–175 (2016).

27. Krill-Burger, J. M. et al. Renal Cell Neoplasms Contain Shared Tumor Type–Specific Copy Number Variations. Am. J. Pathol. 180, 2427–2439 (2012).

28. Han, W. K., Bailly, V., Abichandani, R., Thadhani, R. & Bonventre, J. V. Kidney Injury Molecule-1 (KIM-1): A novel biomarker for human renal proximal tubule injury. Kidney Int. 62, 237–244 (2002).

29. Berry, W. L. & Janknecht, R. KDM4/JMJD2 histone demethylases: epigenetic regulators in cancer cells. Cancer Res. 73, 2936–2942 (2013).

30. Cheung, N. et al. Targeting Aberrant Epigenetic Networks Mediated by PRMT1 and KDM4C in Acute Myeloid Leukemia. Cancer Cell 29, 32–48 (2016).

31. Sato, F., Bhawal, U. K., Yoshimura, T. & Muragaki, Y. DEC1 and DEC2 Crosstalk between Circadian Rhythm and Tumor Progression. J. Cancer 7, 153–159 (2016).

32. Ivanova, A., Liao, S.-Y., Lerman, M. I., Ivanov, S. & Stanbridge, E. J. STRA13 expression and subcellular localisation in normal and tumour tissues: implications for use as a diagnostic and differentiation marker. J. Med. Genet. 42, 565–576 (2005).

33. Telliam, G. et al. Generation of an induced pluripotent stem cell line from a patient with chronic myeloid leukemia (CML) resistant to targeted therapies. Stem Cell Res. 17, 235–237 (2016).

34. Gao, J. et al. Integrative analysis of complex cancer genomics and clinical profiles using the cBioPortal. Sc¡. Signal. 6, pl1 (2013).

35. Irizarry, R. A. et al. Summaries of Affymetrix GeneChip probe level data. Nucleic Acids Res. 31, e15 (2003).

36. Breitling, R., Armengaud, P., Amtmann, A. & Herzyk, P. Rank products: a simple, yet powerful, new method to detect differentially regulated genes in replicated microarray experiments. FEBS Lett. 573, 83–92 (2004).

37. Culhane, A. C., Thioulouse, J., Perrière, G. & Higgins, D. G. MADE4: an R package for multivariate analysis of gene expression data. Bioinforma. Oxf. Engl. 21, 2789–2790 (2005).

38. Tibshirani, R., Hastie, T., Narasimhan, B. & Chu, G. Diagnosis of multiple cancer types by shrunken centroids of gene expression. Proc. Natl. Acad. Sci. U. S. A. 99, 6567–6572 (2002).

39. Xia, J., Gill, E. E. & Hancock, R. E. W. NetworkAnalyst for statistical, visual and network–based meta-analysis of gene expression data. Nat. Protoc. 10, 823–844 (2015).

40. Breuer, K. et al. InnateDB: systems biology of innate immunity and beyond—recent updates and continuing curation. Nucleic Acids Res. 41, D1228–D1233 (2013).

41. Ogata, H. et al. KEGG: Kyoto Encyclopedia of Genes and Genomes. Nucleic Acids Res. 27, 29–34 (1999).

42. Subramanian, A. et al. Gene set enrichment analysis: a knowledge-based approach for interpreting genome-wide expression profiles. Proc. Natl. Acad. Sci. U. S. A. 102, 15545–15550 (2005).

